# Dual effect of α-synuclein disease variants on condensate formation

**DOI:** 10.1101/2025.06.06.657340

**Authors:** Aswathy Chandran, Aishwarya Agarwal, Tianhao Wang, Leslie Amaral, Susana R. Chaves, Tiago F. Outeiro, Janin Lautenschläger

## Abstract

α-synuclein is a pre-synaptic protein^1,2^ implicated in synucleinopathies like Parkinson’s disease and Dementia with Lewy Bodies, where it accumulates in intracellular aggregates termed Lewy bodies and Lewy neurites^3^. Recent studies have reported that α-synuclein undergoes phase separation to form biomolecular condensates both *in vitro* and in mammalian cells^4–7^. α-synuclein condensates are thought to contribute to disease through progressive aggregation^5,8,9^. Here we show that specific PD-associated α-synuclein variants fail to form biomolecular condensates. We demonstrate that only two α-synuclein variants associated with familial disease, E46K and E83Q, enhance condensate formation in vitro and in cells. However, variants including A30G, G51D, and A53E fail to form or have reduced levels of condensate formation in cells. The same phenotypes are reflected in the budding yeast model showing differential inclusion formation. In iPSC-derived neurons, the propensity of α-synuclein variants to undergo VAMP2-mediated phase separation reflects the level of synaptic enrichment. We show that the intrinsic propensity of α-synuclein to form condensates and the ability to bind lipid membranes are important to mediate condensate formation in cells. Our results emphasize that α-synuclein pathology follows divergent pathways, with both increased and decreased condensate formation contributing to disease. This study establishes biomolecular condensates as a key intermediate in α-synuclein dysfunction, providing a novel foundation for translational research.

## INTRODUCTION

Parkinson’s disease (PD) is the second most common neurodegenerative disorder, affecting millions worldwide^10,11^. The disease is characterized by cardinal motor symptoms of tremor, rigidity, bradykinesia, and postural instability, as well as non-motor symptoms, including autonomic dysfunction, dementia, and depression^12^. The pathophysiology of PD is multifactorial, involving genetic, environmental, and age-related factors. In PD and other synucleinopathies like Dementia with Lewy bodies (DLB), α-synuclein (aSYN) misfolds and aggregates in neurons, forming β-sheet rich fibrillar intracellular inclusions known as Lewy bodies and Lewy neurites, hallmark features of the disease^3^. Additionally, duplications/multiplications and mutations of the gene encoding aSYN (*SNCA*) cause autosomal dominant forms of early onset PD^13,14^. aSYN is a key presynaptic protein^1,2^ involved in synaptic vesicle clustering, neurotransmitter release, and maintaining synapse integrity^15^. Recent research highlights that aSYN can undergo phase separation, forming protein-enriched droplet-like structures termed biomolecular condensates^4–7^. This mechanism allows proteins to enrich in dynamic structures, which can form, but also disassemble upon physiological cues^16^. aSYN phase separation is modulated by vesicle-associated membrane protein 2 (VAMP2), a SNARE complex protein responsible for synaptic vesicle fusion^6,7^. This process is modulated by the juxtamembrane domain of VAMP2 and aSYN condensates exhibit liquid-like properties^6,7^.

In this study, we explore the effect of disease-associated aSYN variants on its phase separation *in vitro* and in different cellular environments. We find that only two aSYN variants, E46K and E83Q enhance condensate formation, while certain disease variants show no or markedly reduced condensate formation in cells. aSYN phase separation has been linked to subsequent aggregation and disease pathogenesis^5,8,9^ but our findings highlight that not only increased condensate formation, but an imbalance in aSYN phase separation may contribute to its pathophysiology. This study establishes phase separation as an important molecular property related to aSYN pathology, wherein both, increased or decreased levels of condensate formation, can result in a dysfunction and redistribution of aSYN at the synapse, eventually contributing to disease.

## RESULTS

### aSYN disease variants promote condensate formation

Our previous work showed that aSYN phase separation is regulated by interaction with VAMP2^6^. Using this model of VAMP2-mediated aSYN phase separation in HeLa cells, we now investigate the phase separation behaviour of disease-associated aSYN variants. We find that aSYN variants E46K, A53T, and the newly described DLB-associated variant, E83Q^17^ undergo VAMP2-mediated condensate formation (Fig. 1a). Condensate formation is enhanced for E46K and E83Q, while A53T shows similar levels of condensate formation compared to WT (Fig. 1b). Notably, similar patterns of phase separation are seen *in vitro*, where E46K and E83Q show increased droplet formation, while A53T shows comparable levels of droplet formation to WT (Fig. 1c/d/e). Since phosphorylation of aSYN at serine 129 (pS129) is a well-established pathogenic marker in synucleinopathies, with increased levels seen in Lewy bodies^18,19^ and in response to fibril-induced aggregation in neuronal models^20^, we next examined the phosphorylation status of aSYN condensates in cells. Condensates were consistently pS129 positive, with a significant increase observed in the high-condensate-forming variants, E46K and E83Q. Interestingly, A53T, which is highly aggregation prone^21–23^, exhibits comparable pS129 staining to WT (Extended Data Fig. 1a/b). We further tested whether small aliphatic alcohols, such as 1,6-hexanediol, influence condensate stability in cells^24^. All aSYN condensates, regardless of the variant, dispersed upon treatment with 1,6-hexanediol (Extended Data Fig. 1c). Fluorescence recovery after photobleaching (FRAP) analysis further confirms that aSYN condensates, regardless of the mutation, exhibit rapid recovery kinetics, indicating a high degree of mobility between the condensates and the cytosolic aSYN pool (Extended Data Fig. 1d).

**Figure 1.**
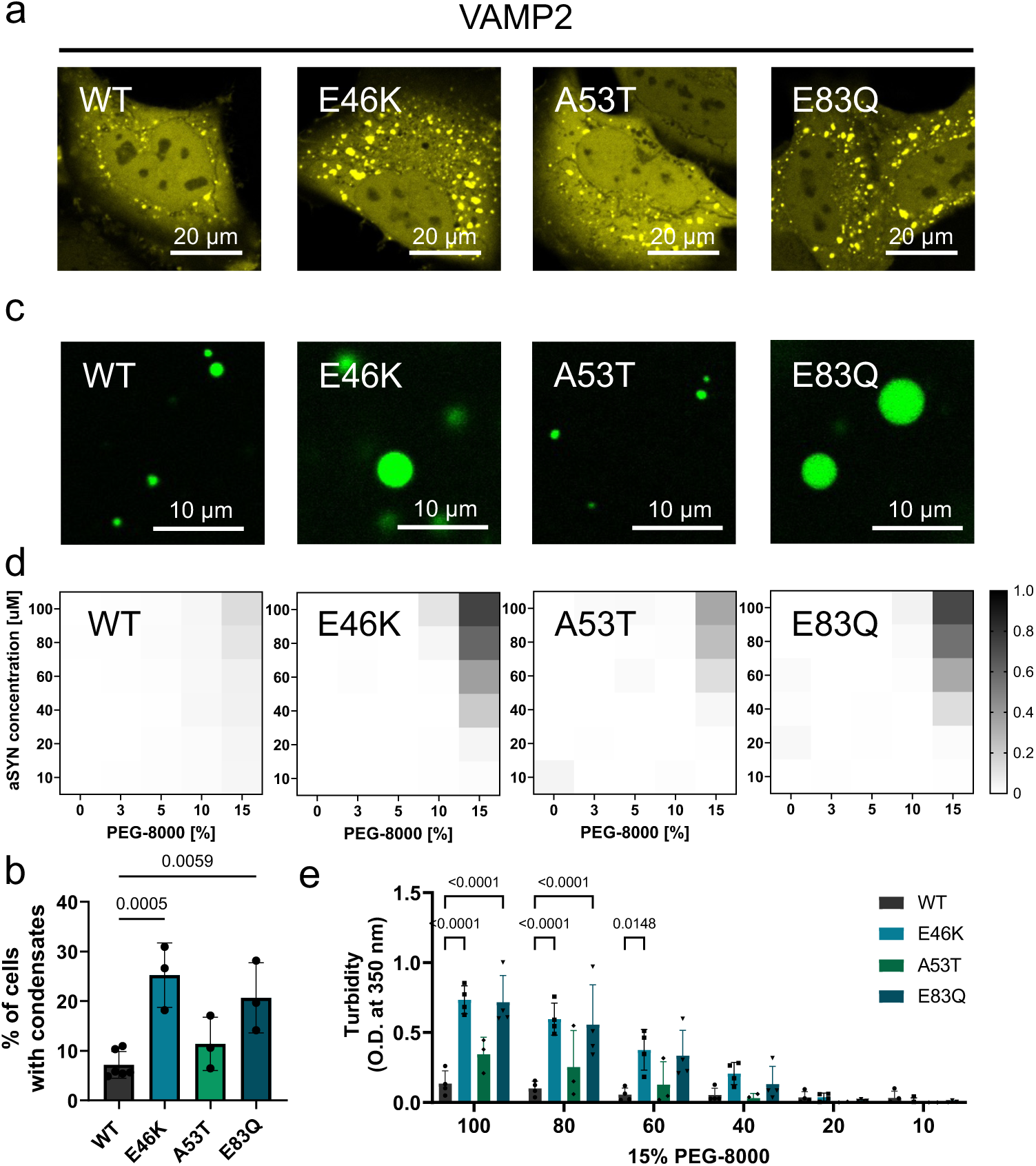
aSYN disease variants promote condensate formation. a, Condensate formation of aSYN-YFP upon co-expression with VAMP2 in HeLa cells. Increased condensate formation is observed for E46K-YFP and E83Q-YFP. b, Quantification of condensate formation in cells, data derived from Incucyte screening with 16 images per well, three wells per biological repeat and at least three biological repeats; n indicates biological repeats. Data are shown as mean ± s.d. One-way ANOVA with Dunnett’s multiple comparison test. c, aSYN phase separation in the presence of 2 mM Ca^2+^ and crowding with 15% PEG 8000, αSYN was used at 100 µM, supplemented with 1% aSYN-Alexa 488, 25 mM HEPES buffer pH 7.4. d, Quantification of aSYN phase separation *in vitro* using turbidity measurements shown as heatmaps for WT, E46K, A53T, and E83Q. Data represent at least 3 independent repeats from two aSYN batches. e, Quantification of turbidity data in d. n represents independent repeats from two aSYN batches. Data are shown as mean ± s.d. Two-way ANOVA with Tukey’s multiple comparison test.

### aSYN disease variants abolish condensate formation

In our previous study, we reported that A30P fails to form condensates in cells^6^. Here we observe that other aSYN variants, including A30G and G51D also show no condensate formation in HeLa cells (Fig. 2a/b). Additionally, A53E has markedly reduced levels of condensate formation, however a small subset of cells form condensates (Fig. 2a/b). We next examined whether these variants would form condensates *in vitro*. All variants were capable of undergoing phase separation *in vitro*, albeit to a lesser extent for G51D and A53E (Fig. 2c/d, Extended Data Fig. 2a). A30P and A30G exhibit similar or slightly higher phase separation compared to WT, in contrast to our observations in cells. Since A30P is known to have reduced lipid binding properties^25^, we next compared lipid binding of the other variants using circular dichroism experiments where aSYN shows a transition from random coil to α-helix in the presence of increasing amounts of small unilamellar vesicles (SUVs, Extended Data Fig. 2b). A30P shows reduced α-helical content in the presence of SUVs, consistent with impaired lipid binding. To a lesser degree, A30G, G51D, and A53E also show reduced α-helical content (Extended Data Fig. 2c). Taken together, these results suggest that, for A30P and A30G, an absence of condensate formation in cells cannot be explained by their intrinsic phase separation propensity, but is likely to be affected by reduced lipid binding. For G51D and A53E, lack of condensate formation in cells could result from a combination of both factors, decreased intrinsic phase separation propensity, but also lipid binding deficiency.

**Figure 2.**
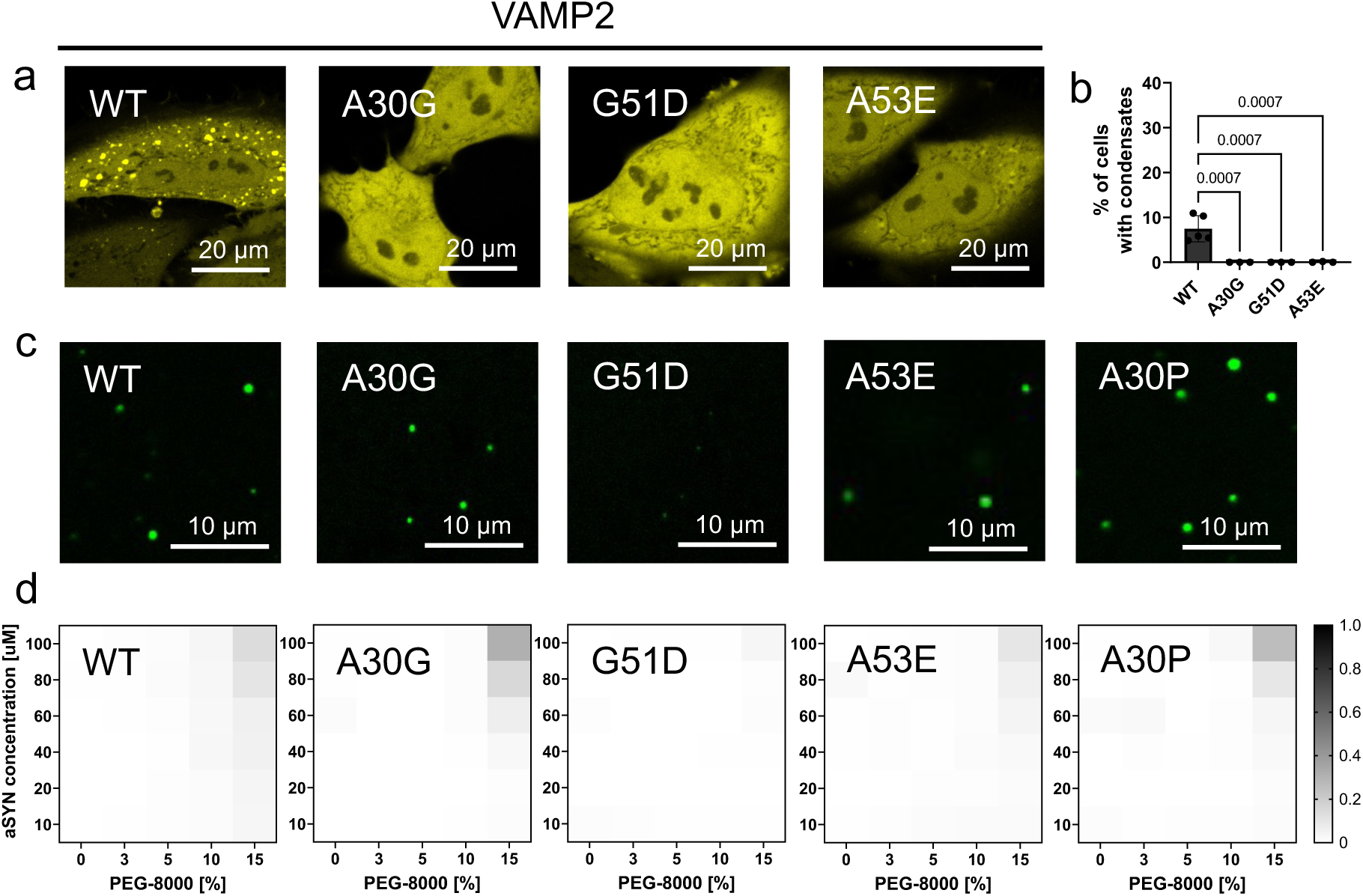
aSYN disease variants reduce condensate formation. a, Condensate formation of aSYN-YFP upon co-expression with VAMP2 in HeLa cells. No condensate formation is observed for A30G and G51D, only very small condensates are seen in a subset of cells for A53E-YFP. b, Quantification of condensate formation in cells, data derived from Incucyte screening with 16 images per well, three wells per biological repeat and at least three biological repeats; n indicates biological repeats. Data are shown as mean ± s.d. One-way ANOVA with Dunnett’s multiple comparison test. c, Recombinant aSYN phase separation *in vitro* in the presence of 2 mM Ca^2+^ and crowding with 15% PEG 8000, αSYN was used at 100 µM, supplemented with 1% aSYN-Alexa 488, 25 mM HEPES buffer pH 7.4. d, Quantification of aSYN phase separation using turbidity measurements shown as heatmaps for WT, A30G, G51D, A53E and A30P aSYN. Data represent 4 independent repeats from two aSYN batches.

### Lipid binding promotes aSYN phase separation

Our results so far show that aSYN variants can have contrasting effects on VAMP2-mediated condensate formation in cells; where some variants facilitate but others abolish condensate formation. To further confirm that the observed phenotypes for the different variants translate to other model systems, we expressed aSYN in budding yeast, which has been established as a model for a distinct cellular environment in which aSYN displays membrane enrichment and inclusion formation^26^. Strikingly, we find that the pattern of inclusion formation of aSYN variants reflects the phenotypes observed upon VAMP2-mediated aSYN condensate formation in HeLa cells. While WT, E46K and A53T form inclusions in yeast, A30P, A30G, G51D and A53E show no inclusion formation (Fig.3a). A53E, however, shows plasma membrane localization, which may demonstrate a pre-inclusion state and might relate to its intermediate condensate phenotype in HeLa cells. Due to the potential role of lipid membranes on aSYN condensate formation, we next tested whether lipid membranes can affect aSYN phase separation in *in vitro* assays. We first confirmed that aSYN condensates formed *in vitro* incorporate SUVs (Fig. 3b), consistent with our previous finding that aSYN condensates in cells cluster intracellular vesicles^6^. We further find that SUVs can increase aSYN droplet formation *in vitro* (Fig. 3c/d), which is also seen in quantitative turbidity measurements (Fig. 3e). Since SUVs are too small to be imaged individually, we turned to giant unilamellar vesicles (GUVs) as another model system. When aSYN, labelled with Alexa 488, was incubated with GUVs, aSYN was uniformly distributed outside of GUVs, with no enrichment on the surface (Fig. 3f). However, when condensate formation was subsequently induced by addition of Ca^2+^ and PEG to the aSYN-GUV mixture, we see aSYN condensates preferentially form on the GUV surface (Fig. 3g). If GUVs were added after induction of phase separation with Ca^2+^ and PEG, aSYN droplets mostly remained in solution, showing significantly less co-localization with GUVs (Fig. 3h/i). This suggests that aSYN condensates form on the GUVs rather than simply binding to their surface, supporting the concept that aSYN condensates can nucleate on lipid membranes.

**Figure 3.**
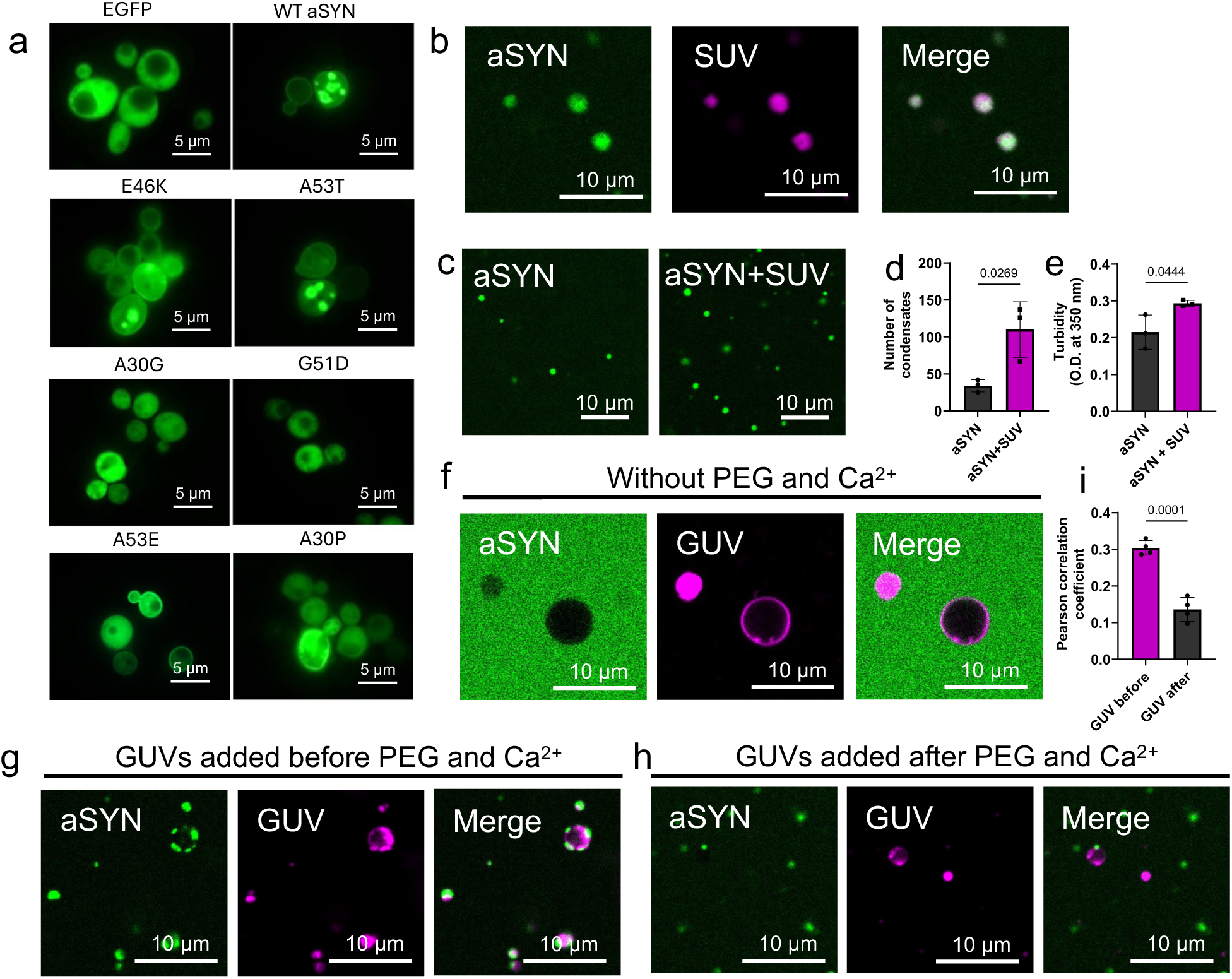
aSYN inclusion formation in yeast and role of lipid membranes. a, aSYN inclusion formation in yeast showing EGFP as a control with cytosolic distribution, while aSYN-EGFP WT forms inclusions. Inclusion formation is also seen for E46K-EGFP and A53T-EGFP. A30G-EGFP, G51D-EGFP, and A30P-EGFP show a cytosolic distribution, while A53E-EGFP shows cytosolic distribution and plasma membrane enrichment. b, Recombinant aSYN phase separation in the presence of 1 mM SUVs, 2 mM Ca^2+^ and crowding with 15% PEG 8000. αSYN was used at 80 µM, supplemented with 1% aSYN-Alexa 488, in 25 mM HEPES buffer pH 7.4. SUV were labelled containing 0.1% PE-Cy5. c, aSYN phase separation with 2 mM calcium and 15% PEG 8000, without and with 1 mM SUVs, 25 mM HEPES buffer pH 7.4. d, Quantification of number of condensates from imaging in c. n indicates biological repeats. Data are shown as mean ± s.d. Unpaired two-tailed t-test. e, Quantification of aSYN phase separation without and with 1 mM SUVs using turbidity measurements. n indicates biological repeats. Data are shown as mean ± s.d. Unpaired two-tailed t-test. All SUVs containing 60% DOPC, 30% DOPS, and 10% cholesterol. f, aSYN incubated with 1 mM GUVs. αSYN was used at 100 µM, supplemented with 1% aSYN-Alexa 488. GUVs labelled containing 0.1% PE-Cy5. g/h, aSYN was incubated in the presence of 2 mM calcium and 15% PEG 8000 to induce aSYN phase separation. GUVs were added either before αSYN phase separation was induced or after aSYN phase separation was induced. aSYN was used at 100 µM, supplemented with 1% aSYN-Alexa 488. GUV labelled containing 0.1% PE-Cy5. GUVs containing 60% DOPC, 30% DOPS, and 10% cholesterol. i, Analysis of aSYN and GUV co-localisation using Person correlation. n indicates biological repeats. Data are shown as mean ± s.d. Unpaired two-tailed t-test.

We returned to our HeLa cell model to determine whether lipid membranes are necessary for VAMP2-mediated aSYN condensate formation. To this end, we expressed a truncated variant of VAMP2 (VAMP2 1-96), which lacks the C-terminal transmembrane domain. VAMP2 1-96 localized to the cytosol (Fig. 4a), and, when co-expressed with aSYN-YFP, was not able to induce aSYN condensate formation (Fig. 4b/c). Full-length VAMP2 however, localizes to vesicular structures and ER membranes upon overexpression (Fig. 4d). We therefore investigated the co-localisation of aSYN condensates with organelle markers, including lysosomes, ER and mitochondria (Extended Data Fig. 3a). Here we find that aSYN condensates mainly reside on the ER tubule network (Extended Data Fig. 3a), consistent with the enrichment of VAMP2 on ER membranes in our model system. Using three-color live-cell high-resolution Airyscan imaging, we observe that aSYN condensates are associated with VAMP2-positive ER membranes and time resolved imaging demonstrates that they move on the ER network showing fluid-like properties (Extended Data Fig. 3b, Fig. 3e, Movie 1 and 2). Together, these findings further support the concept that aSYN lipid binding is necessary for its condensate formation.

**Figure 4.**
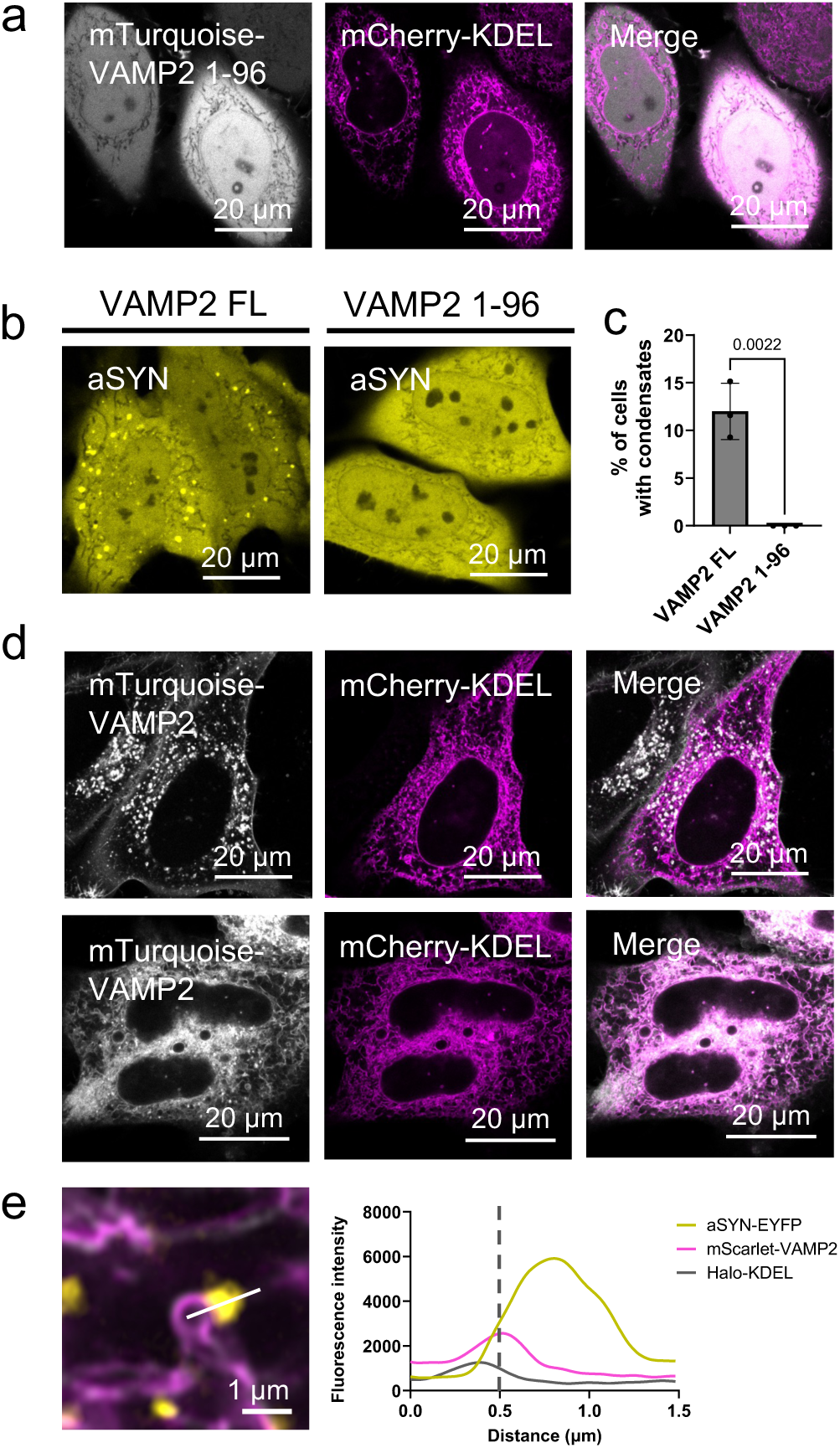
aSYN condensate formation on lipid membranes in cells. a, HeLa cells co-expressing mTurquoise VAMP2 1-96 show cytosolic distribution of VAMP2 lacking its transmembrane domain, mCherry-KDEL shows ER labelling. b, HeLa cells co-expressing aSYN-YFP WT with VAMP2 full length (FL) show aSYN condensate formation, HeLa cells co-expressing aSYN-YFP WT with VAMP2 1-96 show no condensate formation. c, Quantification of condensate formation in cells, data derived from Incucyte screening with 16 images per well, three wells per biological repeat and three biological repeats; n indicates biological repeats. Data are shown as mean ± s.d. One-way ANOVA with Dunnett’s multiple comparison test. d, HeLa cells expressing mTurquoise-VAMP2 FL show a mixed localization pattern with both vesicular and ER distribution, mCherry-KDEL is used to visualize ER network. e, HeLa cells co-expressing aSYN-YFP, Halo-KDEL, and mScarlet-VAMP2 FL show aSYN condensates on VAMP2 positive ER membranes, also see Extended Data Figure 3 for organelle co-localisation and Supplementary Movie 1 and 2.

### Synaptic enrichment of aSYN disease variants

To examine whether aSYN variants may have different patterns of cellular localization in neurons, we utilized an iPSC-derived neuron model to determine levels of synaptic enrichment of aSYN. CRISPR-Cas9 gene editing was used to knock out endogenous *SNCA* in an iPSC line with stable NGN2 overexpression, followed by differentiation to cortical neurons^27,28^ (for characterisation, see Supplementary Fig. 1). Lentiviruses were used to transduce neurons with WT aSYN or disease-variants tagged to the fluorescent protein mScarlet to enable live cell imaging. WT aSYN-mScarlet shows enriched localization at the synapse, although a cytosolic fraction is also present. mScarlet on its own shows no synapse enrichment with an evenly cytosolic distribution (Fig 5a). We proceeded to investigate the distribution of the aSYN disease variants in neurons. Similar to WT aSYN, E46K, A53T and E83Q variants also show synaptic enrichment (Fig. 5b). However, variants that do not undergo VAMP2-mediated phase separation in HeLa cells, fail to enrich at synaptic terminals in neurons. This is the case for A30G, G51D and A30P, which have an evenly distributed cytosolic expression, similar to mScarlet alone (Fig. 5c). A53E also exhibits a predominant cytosolic expression, however we observe synaptic enrichment in a subset of terminals (Fig. 5c/d). We quantified the intensity ratio of aSYN at synapses using a synaptophysin mask. This confirms synaptic enrichment of WT, E46K, A53T and E83Q, while for A30G, G51D, A53E and A30P synaptic localization is significantly reduced (Fig 5e). We further quantified the number and size of aSYN enriched synapses for WT, E46K, A53T and E83Q, wherein we see no changes for the number of aSYN-enriched synapses (Fig. 5f) but an increase in size for E46K and E83Q compared to WT (Fig. 5g). No increase in the size of aSYN-enriched synapses is seen for A53T, in line with our in vitro and HeLa cell observations where A53T behaves similarly to WT. To confirm that reduced synaptic enrichment is not caused due to lower protein expression of the respective variants, we performed western blotting demonstrating that there are no significant differences between expression levels among the variants, suggesting that differential expression levels are not the cause of increased or decreased synaptic localization (Extended data Fig. 4).

**Figure 5.**
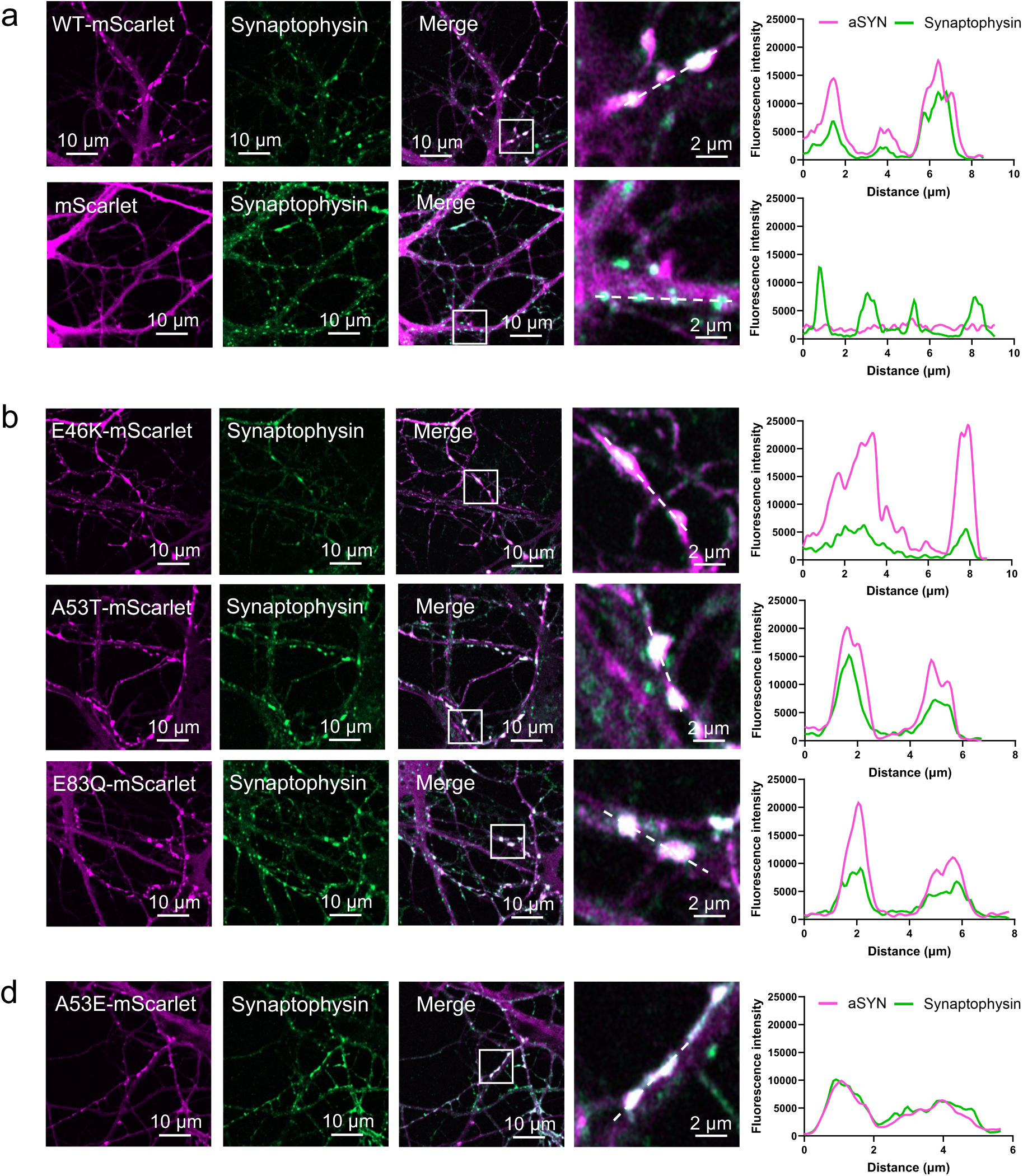
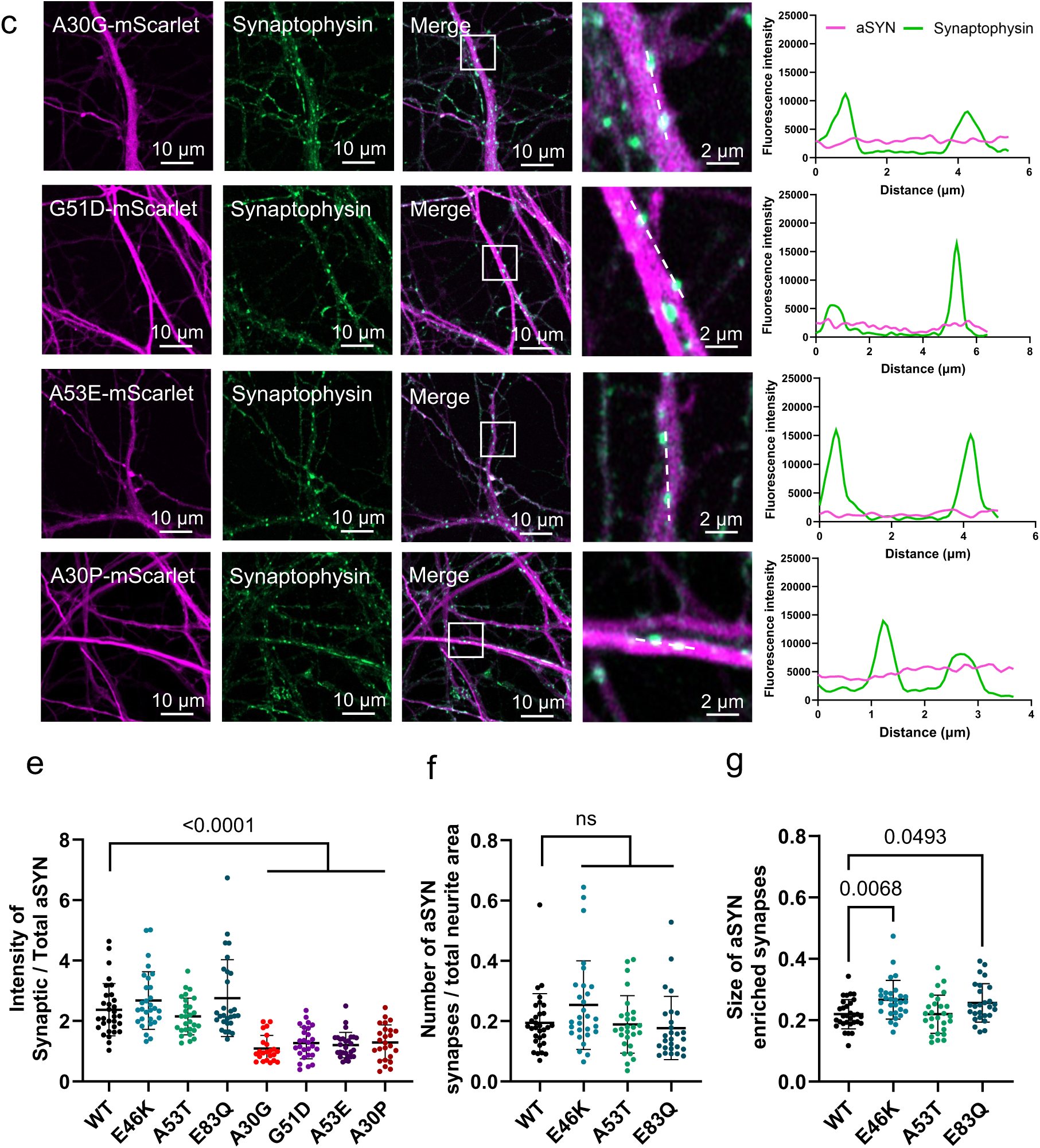
Synaptic enrichment of aSYN disease variants. a, WT aSYN-mScarlet and mScarlet expressed in SNCA-knock out iPSC-derived cortical neurons. iPSC-derived neurons were transduced with lentiviruses between D21-25 and imaged five days post transduction. Synaptophysin-mEmerald was expressed as synapse marker. Zoomed in images and line diagrams for synapse enrichment are indicated. b, E46K-, A53T- and E83Q-mScarlet expressed in SNCA-knock out iPSC-derived cortical neurons. Zoomed in images and line diagrams showing synapse enrichment. c, A30G-, G51D-, A53E- and A30P-mScarlet expressed in SNCA-knock out iPSC-derived cortical neurons. Zoomed in images and line diagrams showing missing synapse enrichment. d, Example images of A53E-mScarlet showing synaptic enrichment in certain terminals. Zoomed in image and line diagram showing synapse enrichment. e, Quantification of aSYN synapse enrichment displayed as a ratio of intensity of aSYN at the synapse vs. total aSYN. Synaptic terminals were selected using the synaptophysin mask. n indicates number of fields of view, n= 31, 29, 28, 28, 23, 27, 28 and 25 for WT, E46K, A53T, E83Q, A30G, G51D, A53E and A30P respectively. Data from 3 biological repeats. Data shown as mean ± s.d. One-way ANOVA with Dunnett’s multiple comparison test. f, Quantification of the number of aSYN-enriched synapses normalized to the neurite area. Cell bodies were excluded from area measurements. No significant differences are seen between WT, E46K, A53T and E83Q expressing neurons. g, Quantification of the size of aSYN-enriched synapses in WT, E46K, A53T and E83Q expressing neurons showing an increase in size for E46K and E83Q compared to WT. n indicates number of fields of view with n=31, 29, 27, 27 for WT, E46K, A53T, and E83Q respectively. Data from 3 biological repeats. Data are shown as mean ± s.d. One-way ANOVA with Dunnett’s multiple comparison test.

As demonstrated for other soluble phase separating proteins, such as dynamin^29^ and endophilin A1^30^, we tested whether synaptic enrichment of aSYN is sensitive to 1,6-hexanediol. We see that aSYN rapidly disperses from synapses upon incubation with 1,6-hexanediol and furthermore, we observe a rapid recovery of synaptic aSYN enrichment upon 1,6-hexanediol washout (Fig. 6a/b). To characterise the behaviour of aSYN at synapses further, we performed FRAP experiments comparing aSYN fluorescence recovery at the synapse and in dendrites. We observe a clearly distinct recovery profile for synaptic aSYN, showing significantly slower recovery in synapses compared to dendrites, demonstrating that aSYN is actively retained in synapses and not freely diffusive (Fig. 6c). We subsequently performed FRAP experiments at the synapse for all synaptically enriched aSYN variants. We find that all aSYN variants retain high mobility at the synapse, consistent with the fluorescence recovery profiles seen for aSYN condensates in HeLa cells. Only E46K shows a small decrease in the extent of recovery (Fig. 6d). The link between condensate formation of aSYN disease variants in HeLa cells and their synaptic enrichment in neurons suggests that phase separation might be important to regulate aSYN subcellular compartmentalisation. Taken together, our findings demonstrate that while aSYN condensate formation is increased for some disease variants like E46K and E83Q, this is not the case for other variants, including A30G, G51D, A53E and A30P, indicating that a disturbance of condensate formation, both increased as well as decreased, may contribute to disease.

**Figure 6.**
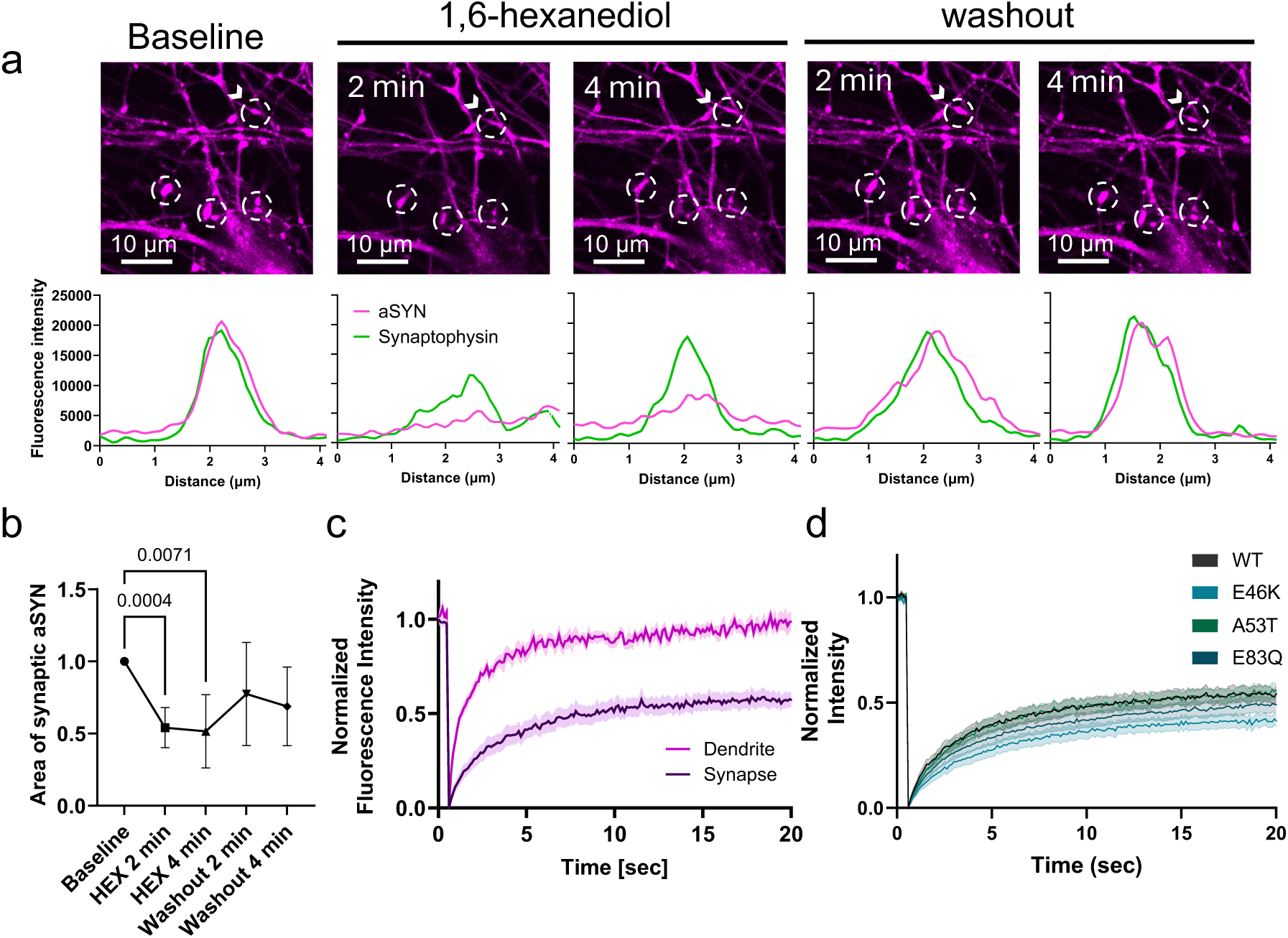
Characterisation of aSYN synapse enrichment. a, WT aSYN-mScarlet expressed in SNCA-knock out iPSC-derived cortical neurons showing synapse enrichment at baseline. Neurons were incubated with 3% 1,6-hexanediol for 2 and 4 min, then 1,6-hexanediol was washed out and new media added. Synaptophysin-mEmerald was expressed as synapse marker. b, Quantification of aSYN synaptic enrichment before, upon incubation with 3% 1,6-hexanediol and upon washout. n indicates number of fields of view with 7 images analysed from 3 biological repeats. Data are shown as mean ± s.d. RM one-way ANOVA with Geisser-Greenhouse correction and Dunnett’s multiple comparisons test. c, Fluorescence recovery after photobleaching (FRAP) for aSYN-mScarlet in dendrites and at synapses, showing faster and greater extent of recovery in dendrites compared to synapses. Data are shown as mean ± s.e.m. n indicates number of individual FRAP measurements, n=9 for both synaptic and dendritic FRAP from 3 biological repeats. d, FRAP for WT-, E46K-, A53T- and E83Q-mScarlet at synapses. Data are shown as mean ± s.e.m., n indicates number of synapses measured, with n=26, 22, 25 and 27 for WT, E46K, A53T, and E83Q respectively. Data from 3 biological repeats.

## DISCUSSION

In summary, we find that disease associated aSYN variants exhibit a dichotomy in their ability to undergo biomolecular condensation, wherein only E46K and E83Q demonstrate enhanced condensate formation both in cells and *in vitro*. In contrast, A30P, A30G, G51D and A53E form no or very few condensates in cells. Our findings related to E46K are consistent with previously published work which has demonstrated that E46K has increased phase separation propensity in vitro^4,31^. Enhanced phase separation of E83Q, a variant associated with dementia with Lewy bodies^17,32^, has not been described before. Interestingly, A53T, the most widely studied aSYN disease variant which is directly associated with increased aggregation^21,22,33^, displays phase separation comparable to aSYN WT^4,31^. We show that all condensates exhibit pS129 immunoreactivity in HeLa cells, with E46K and E83Q variants showing greater levels of pS129, while A53T was comparable to WT. Recent studies have called into question the strict association between pS129 and pathological aSYN aggregation, suggesting that pS129 also plays a physiological role^34,35^. Subsequently, aSYN disease variants were also shown to exhibit different tendencies to undergo pS129 phosphorylation under non-aggregating conditions. E46K has been reported to have higher levels of basal pS129 phosphorylation, while A53T exhibited similar levels as WT^36^, consistent with our findings here. Moreover, we also observe dispersion of aSYN condensates upon incubation with 1,6 hexanediol and high fluorescence recovery in FRAP experiments. Collectively, our data therefore suggest that in our HeLa cell model, VAMP2-mediated aSYN condensates, irrespective of the disease-associated mutations, are dynamic structures lacking significant aggregation into insoluble conformations.

Four aSYN variants, A30P, A30G, G51D and A53E show absent or markedly reduced condensate formation in cells. However, in *in vitro* assays both A30P and A30G undergo significant phase separation. This is consistent with previous work, showing that A30P can undergo condensate formation *in vitro* comparable to WT^31,37^ but has not been reported for A30G. Our circular dichroism experiments with A30P indicate reduced alpha-helicity in the presence of lipid vesicles, suggesting reduced lipid binding, consistent with previous studies^25,38,39^. Similarly, but to a lesser extent, A30G also shows reduced alpha-helicity in the presence of lipid vesicles. Therefore, the discrepancy between *in vitro* and cellular condensate formation by A30P and A30G is likely to be explained by their decreased membrane binding characteristics. We therefore tested whether lipid binding can facilitate aSYN phase separation. We show that lipid membranes (SUVs and GUVs) indeed foster aSYN droplet formation *in vitro*, and also that aSYN condensates in cells are localised to membrane surfaces. Importantly, membrane associated phase separation has been described for other proteins, such as RIM and RIM-BP^40^ in association with voltage gated calcium channels and for Whi3^41^ and AGO^42^ proteins on GUVs and in cells, supporting our observations on membrane surfaces acting as a scaffold for aSYN phase separation. For G51D and A53E, we see reduced alpha-helicity in CD experiments, consistent with decreased lipid binding that has been described before^43–45^, however these variants additionally show decreased droplet formation *in vitro*. Therefore, the lack of biomolecular condensate formation by these variants in cells could demonstrate a combined effect of decreased intrinsic phase separation propensity and lowered membrane binding.

Finally, we show that the phenotype of VAMP2-mediated aSYN condensate formation in HeLa cells translates to the propensity of aSYN to enrich at synaptic terminals in neurons, with WT, E46K, A53T and E83Q showing synapse enrichment, while A30P, A30G, G51D and A53E exhibiting a predominant cytosolic distribution. A53E however, shows enrichment in some synaptic terminals, similar to our observations that A53E can form condensates in a subset of HeLa cells. Given that aSYN phase separation is influenced by VAMP2, a synaptic vesicle protein, and the ability of aSYN condensates to cluster vesicles in cells^6^ similar to synapsin 1^46^, we propose that synaptic enrichment of aSYN could reflect condensate formation at the synapse. This is further evidenced by sensitivity of synaptic aSYN to 1,6-hexanediol and differential FRAP kinetics. Lack of synaptic enrichment has been previously described for A30P^47^ and recently also for G51D^48^, however this has not been shown for A30G and A53E, further emphasizing that this phenotype may be directly relevant to disease. With this, we suggest that in addition to aggregation, differential condensate formation by aSYN disease variants may have distinct cellular effects, either via protein redistribution from the synapse to other cellular compartments or via a direct effect on synaptic function, wherein aSYN may lose its ability to participate in vesicle clustering, exocytosis or endocytosis^15^. Similarly, enhanced phase separation may directly affect activity dependent condensation and dispersion of aSYN at the synapse, causing changes in synaptic function, prior to aggregation. Previous studies on aSYN phase separation have linked subsequent liquid-to-solid transition of aSYN condensates *in vitro* to the formation of insoluble fibrils ^4,8,9,49^. While this pathway may still hold true for variants that promote phase separation, we challenge this view as the only mechanism of aSYN pathology associated with phase separation. aSYN variants with decreased condensate formation leads to a redistribution of synaptic aSYN, which could result in its aggregation in the cytosol, similar to what has been recently described for G51D mutant mice wherein fibril induced aggregation was found to be increased, compared to WT mice^48^. Likewise, a redistribution of aSYN from the synapse could also affect other organelle systems. Additionally, such changes in localization may lead to formation of toxic aSYN oligomers which has been reported for these variants^44,50–54^. We further aim to highlight that particularly G51D and A53E variants, which have decreased intrinsic property to phase separate, lead to an early age of onset in people with PD^55–60^, and are therefore highly relevant to be explored further. Taken together, our data demonstrate an unexpected dichotomy associated with the dysregulation of aSYN phase separation, which may directly affect disease pathogenesis.

## METHODS

### Plasmids

Constructs for aSYN-YFP and VAMP2 were used from previous studies^6^. Briefly, wild-type human full-length SNCA and VAMP2, encoding aSYN and VAMP2, were cloned from cDNA obtained from human neuroblastoma cells (SH-SY5Y) and inserted into the pEYFP-N1 and pMD2.G vector (Addgene #96808, #12259) with a C-terminal YFP and Flag-tag, respectively. Gibson assembly was performed upon PCR amplification (Q5 Hot start HiFi 2xMM, M0494S, 2xHiFi DNA Assembly MM, E2621S, NEB, Ipswich, US). aSYN A30P, A30G, E46K, G51D, A53T, A53E and E83Q; and VAMP2 1-96 were generated using KLD substitution (M0554S, NEB, Ipswich, US). 5HT6-YFP-Inpp5e was a gift from Takanari Inoue (Addgene plasmid # 96808; http://n2t.net/addgene:96808; RRID:Addgene_96808)^61^. pMD2.G was a gift from Didier Trono (Addgene plasmid # 12259; http://n2t.net/addgene:12259; RRID:Addgene_12259). 4xmts-mScarlet-I was a gift from Dorus Gadella (Addgene plasmid # 98818; http://n2t.net/addgene:98818; RRID:Addgene_98818)^62^. pLAMP1-mCherry was a gift from Amy Palmer (Addgene plasmid # 45147; http://n2t.net/addgene:45147; RRID:Addgene_45147)^63^. The PrSS-Halo-KDEL plasmid was generated in the Nixon-Abell lab through three-part Gibson assembly of a HaloTag protein with a flanking Prolactin signal sequence and KDEL retention motif into a standard N1 backbone ^64^. mRFP1-N1 was a gift from Robert Campbell & Michael Davidson & Roger Tsien (Addgene plasmid # 54635; http://n2t.net/addgene:54635; RRID:Addgene_54635)^64^. mCherry-KDEL was created from the PrSS-Halo-KDEL plasmid exchanging Halo for mCherry using Gibson Assembly (Q5 Hot start HiFi 2xMM, M0494S, 2xHiFi DNA Assembly MM, E2621S, NEB, Ipswich, US). All organelle plasmids were gifts from Jonathan Nixon-Abell, University of Cambridge. pmD2G-mTurquoise-VAMP2-FLAG and pmD2G-mScarlet-VAMP2-FLAG plasmids were created inserting mTurquoise2 or mScarlet into the pmD2G VAMP2 Flag plasmid using Gibson Assembly (Q5 Hot start HiFi 2xMM, M0494S, 2xHiFi DNA Assembly MM, E2621S, NEB, Ipswich, US). mTurquoise2 and mScarlet-I were amplified from plasmids ITPKA-mTurquoise2 (Addgene plasmid # 137811; http://n2t.net/addgene:137811; RRID:Addgene_137811) and pmScarlet_Giantin_C1, (Addgene plasmid # 85048; http://n2t.net/addgene:85048; RRID:Addgene_85048) both gifts from Dorus Gadella^62,65^. pmD2G-mTurquoise2-VAMP2_1-96-FLAG was derived from pmD2G-mTurquoise-VAMP2-FLAG plasmid through KLD deletion (M0554S, NEB, Ipswich, US). All plasmids were verified by sequencing.

### Cell culture and transfection

HeLa cells were obtained from the European Collection of Cell Cultures (ECACC 93021013) and grown in Dulbecco’s modified Eagle’s Medium (DMEM) high glucose (31966-021, Gibco) supplemented with 10% fetal bovine serum (FBS, F7524, Sigma), 20 mM HEPES (Sigma, H0887) and 1% Penicillin/Streptomycin (P0781, Sigma). Cells were grown at 37 °C in a humidified incubator with 5% CO2. Cells were tested for mycoplasma contamination using MycoStripTM (IvivoGen, Toulouse, France). Cells were plated at 10,000 cells/well in 8-well ibidi dishes (80807, ibidi, Gräfelfing, Germany) for confocal imaging or 48-well plates (Cellstar, 677 180, Greiner bio-one) for incuCyte experiments. Cells were transfected the following day using Fugene HD Transfection reagent according to the manufacturer’s protocol (E2311, Promega). Briefly, per reaction 12.5 μL OptiMEM (31985-062, Gibco) were set up in 1.5 mL sterile Eppendorf tubes. A total of 250 ng of DNA and 0.75 μL of Fugene reagent were added and incubated for 15 min at room temperature. The transfection mix was added to the cells for 1 min and then topped up with 300 μL complete media. Cells were imaged the next day.

### Microscopy, IncuCyte, and image analysis

Confocal imaging of condensate formation in HeLa cells was performed on an LSM780 microscope (Zeiss, Oberkochen, Germany) using a 63x oil immersion objective. YFP fluorescence was excited with the 514 nm laser. To evaluate aSYN condensate dispersion 1,6-hexanediol (240117, Sigma, USA) was prepared as 6% stock solution in complete DMEM media and added to the cells in a 1:1 ratio after the first image was taken. For fluorescence recovery after photobleaching (FRAP) experiments, images were taken with the 63x oil immersion objective, 20x zoom, 128×128 pixel resolution, at an imaging speed of 60 ms/image. Three pre-bleach images were acquired before the ROI was bleached with 100 iterations at 100% laser power using the 514 nm laser. Fluorescence recovery was recorded for 100 cycles. FRAP analysis was performed in FIJI using the FRAP profiler v2 plugin (Hardin lab, https://worms.zoology.wisc.edu/research/4d/4d.html). Organelle co-localisation studies in HeLa cells were performed on an LSM 780 or LSM880 microscope (Zeiss, Oberkochen, Germany) using a 63x oil immersion objective and Super-resolution (SR) Airyscan processing. YFP was excited with the 488 nm laser, mTurquoise was excited with the 405 nm laser, mCherry and mScarlet were excited using the 561 nm laser. For experiments involving Halo-KDEL, Janelia Fluor® 646 HaloTag® Ligand (GA1120, Promega) was added to the cells at the time of transfection at 1:10000 dilution. HaloTag treated cells were subsequently imaged with the 633 nm laser. Time course imaging was performed using the Super-resolution (SR) Airyscan feature of the LSM 880 microscope. Profile plots were generated in FIJI^66^ and plotted using GraphPad Prism 10.4.1. Zen 2.3 (black edition) and Zen 2.6 (blue edition) were used for data collection. For quantitative experiments wherein we evaluated % of cells with condensates, Hela cells were plated in 48-well plates and imaged with the IncuCyte S3 (Essen BioScience, Newark, UK). Phase bright field and green fluorescence images were taken using a 20x objective at a 4-hour interval at 200 ms exposure. Condensate formation (% of cells showing condensate formation) was evaluated at 16 hours after transfection. IncuCyte 2021A was used for data analysis. At least three biological repeats with three technical repeats each were analysed blinded to the investigator.

### Immunocytochemistry

The following immunocytochemistry (ICC) protocol was used for both HeLa cells and iPSC-derived neurons. Cells were fixed using 4% paraformaldehyde in phosphate-buffered saline (PBS), pH 7.4 for 30 minutes at room temperature. Blocking and permeabilization were performed using 10% FBS, 1% BSA and 0.3% TritonX-100 in PBS for 1h. Cells were stained with the corresponding primary antibodies (refer to Supplementary Antibodies table) in PBS containing 1% BSA, and incubated overnight at 4 °C. The following day, secondary antibodies at 1:1000 in PBS with 1% BSA was added to the cells and incubated at room temperature for 1 hour. Subsequently, the cells were imaged on an LSM780 or LSM 980 confocal microscope (Zeiss, Oberkochen, Germany). For pS129 staining experiments in HeLa cells, aSYN-YFP was excited using the 514 nm laser and pS129 labelled with secondary antibody Alexa Fluor 594 was excited with the 561 nm laser. The ratio of pS129 to total aSYN intensity in the condensates was measured using FIJI^66^. Briefly, a mask was created for the aSYN-YFP channel to select the condensates, which was then applied to the pS129 channel to obtain fluorescence intensities specifically in the condensates. iPSCs stained for pluripotency markers were treated with Alexa Fluor 488 secondary antibody and subsequently imaged with the 488 nm laser. Finally, endogenous alpha-synuclein and β-tubulin in iPSC neurons were stained with secondary antibodies Alexa Fluor 488 and Alexa Fluor 594 respectively and subsequently imaged with the 488 nm and 561 nm laser on the confocal microscope.

### Protein expression and purification of αSYN

Recombinant human full-length aSYN was expressed in BL21(DE3) competent Escherichia coli (C2527, NEB, Ipswich, US) using vector pET28a (Addgene #178032). aSYN A30P, A30G, E46K, G51D, A53T, A53E and E83Q were generated using KLD substitution (M0554S, NEB, Ipswich, US). Bacteria were cultured in LB media supplemented with 50 μg/mL kanamycin (37 °C, constant shaking at 250 rpm). Expression was induced at an OD600 of 0.8 using 1mM isopropyl β-D-1-thiogalactopyranoside (IPTG) and cultured overnight at 25 °C. Cell pellets were harvested by centrifugation at 4000L×Lg for 30 minutes (AVANTI J-26, Beckman Coulter, USA). aSYN was purified using a protocol previously described^67^. Briefly, the cell pellet was resuspended in lysis buffer (10 mM Tris, 1 mM EDTA, Roche cOmplete EDTA free protease inhibitor cocktail, pH 8). The cells were disrupted using a cell disruptor (Constant Systems, Daventry, UK) and were ultra centrifuged at 4 °C, 186,000L×Lg for 20 minutes (Ti-45 rotor, Optima XPN 90, Beckman Coulter, USA). The supernatant was collected and heated for 20 minutes at 70 °C to precipitate heat-sensitive proteins, followed by ultracentrifugation as above. Streptomycin sulphate (5711, EMD Millipore, Darmstadt, Germany) was added at a final concentration of 10 mg/mL to the supernatant and continuously stirred at 4 °C for 15 minutes to precipitate DNA, followed by ultracentrifugation as above. Ammonium sulphate (434380010, Thermo Scientific) was added at a final concentration of 360 mg/mL to the supernatant and continuously stirred at 4 °C for 30 minutes to precipitate the protein. The precipitated protein was then centrifuged at 500Lx g for 15 min, dissolved in 25 mM Tris, pH 7.7, and dialyzed overnight against the same buffer to remove salts. The protein was purified using ion exchange on a HiTrapTMQ HP 5mL anion exchange column (17115401, Cytiva, Sweden) using gradient elution with 0-1M NaCl in 25 mM Tris, pH 7.7. The collected protein fractions were run on SDS-PAGE and pooled fractions were further purified using size-exclusion chromatography on a HiLoadTM 16/600 SuperdexTM 75 pg column (28989333, Cytiva, Sweden). The fractions were collected, and their purity was confirmed using SDS-PAGE analysis. Protein concentrations were determined by measuring absorbance at 280 nm using an extinction coefficient of 5,600 M−1cm−1. The monomeric protein was frozen in liquid nitrogen and stored in 25 mM HEPES buffer pH 7.4 at −70 °C. pET28a Cdk2ap1CAN was a gift from Lin He (Addgene plasmid # 178032; http://n2t.net/addgene:178032; RRID:Addgene_178032)^68^.

### Protein labelling

Labelling of proteins was performed in bicarbonate buffer (C3041, Sigma) at pH 8 using NHS-ester active fluorescent dyes. aSYN was labelled with AlexaFluor 488 5-SDP ester (A30052, Invitrogen Thermo Fisher). Excess-free dye was removed by buffer exchange using PD10 desalting columns (IP-0107-Z050.0-001, emp BIOTECH, Generon). Labelled protein concentrations were estimated using molar extinction coefficients of the dyes with ε494 nmL=L72,000LM−1cm−1 for Alexa-488 5-SDP ester.

### Phase separation assays - turbidity measurements and confocal imaging

Phase separation assays were performed in 25 mM HEPES, pH 7.4 unless mentioned otherwise. Phase separation was induced by mixing aSYN and PEG 8000 (BP223, Fisher Bioreagent) in the presence of 2 mM calcium chloride (21108, Sigma) as indicated respectively. **Turbidity measurements** - Phase-separated samples were set up as described above. The turbidity of the samples was measured at 350 nm, 25 °C using 96-well Greiner optical bottom plates on a CLARIOstar plate reader (BMG LABTECH, Ortenberg, Germany) under quiescent conditions. CLARIOStar 5.01 was used for data acquisition. A sample volume of 100 μL was used, and readings were taken within 5 minutes of sample preparation. For phase diagrams, raw turbidity data are plotted with background subtraction. Data were obtained from four independent repeats and were plotted using GraphPad Prism 10.4.1. **Confocal Microscopy** - Imaging of aSYN condensates in solution or in the presence of SUV or GUVs was performed on an LSM710, 780, or 880 microscope (Zeiss, Oberkochen, Germany) using a 63x oil immersion objective. Alexa 488 was excited with the 488 nm laser, Cy5 was excited using the 633 nm laser. Images were taken using aSYN supplemented with 1% Alexa 488 labelled aSYN. Lipids were labelled incorporating 0.1% Cy5-PE. Zen 2.3 (black edition) and Zen 2.6 (blue edition) were used for data collection. Image analysis was performed in FIJI^66^. To analyse the number of condensates a mask for the droplets was generated, followed by particle analysis. Pearson correlation coefficients were calculated using the ColocFinder Plugin (https://imagej.nih.gov/ij/plugins/colocalization-finder.html).

### Circular dichroism spectroscopy

Circular dichroism (CD) spectra were recorded on a Jasco J-810 Spectrapolarimeter (Jasco, Pfungstadt, Deutschland). aSYN was buffer exchanged into 20 mM sodium phosphate buffer pH 6.26 and incubated at 0.2 mg/mL protein concentration with an increasing amount of SUVs from 0, 0.1, 0.2, 0.3, 0.5 to 1 mM. SUVs were prepared as outlined below, containing 100% POPS, to enable high aSYN binding. Samples were mixed, briefly incubated and measured in a 1 mm pathlength quartz cuvette (QS High Precision Cell, 110-1-40, Hellma Analytics, Southend on Sea, United Kingdom). Spectra were recorded between 200 to 250 nm, with a 0.1 nm step, 1 nm bandwidth, and a speed of 50 nm/minute, averaged over 10 scans. A shift from random coil into α-helical structure was determined from the ellipticity at 222 nm^69^. Measurements were performed at room temperature.

### Small unilamellar vesicles and giant unilamellar vesicles

10 mM stock solutions of small unilamellar vesicles (SUVs) and giant unilamellar vesicles (GUVs) were prepared using a phospholipid mixture of DOPC, DOPS, cholesterol, and Cy5-PE in a 60:30:10:0.1 molar ratio. For SUV preparation, chloroform solutions of the lipids were combined in the required proportions and dried under a nitrogen stream forming a lipid film. The lipid film was hydrated with 25 mM HEPES buffer, pH 7.4 for 2 hours with intermittent vortexing. For circular dichroism (CD) measurements, the lipids were hydrated in 20 mM sodium phosphate buffer, pH 6.26. The resulting lipid suspensions were then subjected to 10 freeze-thaw cycles, using liquid nitrogen and a preheated water bath at 42°C. SUVs were obtained by 2 min probe sonication, 10 sec on; 15 sec off, 20% amplitude (Sonic Dismembrator, Fisherbrand Model 505, Loughborough, UK). Turbidity measurements with SUVs were carried out with 80 µM aSYN and 1 mM lipids unless mentioned otherwise. Turbidity values were subtracted with the respective SUV control containing 1 mM SUV, 2 mM calcium and 15 % PEG 8000 to correct for calcium-induced aggregation of SUVs. GUVs were prepared using a gel-assisted swelling method as previously described^70^. Briefly, glass slides were coated with 100 µL of a 5% polyvinyl alcohol (PVA) solution and heated at 50°C for 20 minutes to form a uniform PVA film. The required amount of lipids was then applied to the PVA-coated surface and dried under a gentle nitrogen stream. The PVA film was subsequently scraped and the lipid film was hydrated with 25 mM HEPES buffer, pH 7.4 for 2 hours, resulting in the formation of GUVs.

### Yeast experiments

aSYN constructs were generated using site-directed mutagenesis carried out using the QuickChange II Site-Directed Mutagenesis Kit (Agilent Technologies, SC, USA) according to the manufacturer’s protocol in the plasmid backbone p426GPDpr encoding SNCA(WT)-GFP. The plasmid without aSyn, p426GPDpr-GFP was constructed by inserting the GFP coding sequence as a SpeI-XhoI digested PCR product. All constructs were confirmed by DNA sequencing. Yeast transformation was performed by standard lithium acetate method as described previously^71^. Supplementary Table 1 shows all aSYN mutants used in this study. Cells were grown overnight in an orbital shaker, at 30 °C, 180 rpm, in yeast minimal synthetic defined (SD) medium (Takara Bio, Shiga, Japan), supplemented with drop-out mix (Takara Bio, Shiga, Japan) lacking the amino acid uracil (SD-URA) for transformant selection in a ratio of flask volume/medium of 5:1. Intracellular distribution of aSYN was evaluated by fluorescence microscopy, images were acquired using a Zeiss Axio Observer microscope equipped with a 100x oil objective.

### iPSC culture and differentiation to cortical neurons

The GM29370 iPSC line with a stable integration of Tet-On human NGN2 at the AAVS1 locus of the WTC-11 line was obtained from NIGMS Human Genetic Cell Repository at the Coriell Institute for Medical Research. iPSCs were cultured in StemFlex media (Gibco, A3349401) according to manufacturer’s guidelines on Matrigel (Corning, 354230) coated plates. 150 μL Matrigel was mixed with 15 mL DMEM/F12 media (Invitrogen, 10565018) and added to plates, which were then incubated at 37°C for an hour before use. Cells were regularly passaged using versene (Gibco, 15040066) for dissociation of iPSCs as clusters of cells or using accutase (Gibco, A1110501) for single cell suspensions. Chroman 1 (Tocris 7163) was used to inhibit ROCK signalling and maintain viability of iPSCs whenever cells were passaged with accutase. At the start of differentiation (D0), iPSCs were passaged with accutase and plated in Neural Induction media containing DMEM/F12 with Glutamax (Gibco 10565018), 1X MEM-Non Essential Amino Acids (Gibco, 11140050) and 1X N-2 supplement (Gibco, 17502048). NGN2 expression was induced by adding Doxycycline (DOX) at 2 μg/ml. On D1 and D2, fresh Neural Induction media supplemented with DOX was added to the cells. On D3, cells were split with accutase and resuspended in complete Neuronal medium containing Neurobasal Plus, supplemented with 1X B27 plus (Gibco A3653401), BDNF 10 ng/mL (Gibco, AF-450-02-10UG), NT-3 10 ng/mL (Gibco, 450-03-10UG), Laminin 1 μg/mL (Sigma, L2020-1MG), DOX 2 μg/mL and the mitotic inhibitor cytosine-β-D-arabinofuranoside/Ara-C 1 μM (Sigma C6645). The cell suspension was plated on 0.01% poly-L ornithine (Sigma, P4957) coated dishes (37°C, overnight) at a density of 12,500 cells per well in an 8-well IBIDI dish (IBIDI, 80807) for live-cell imaging or 375,000 cells per well in a 12 well plate for western blot experiments. The neurons were subsequently allowed to mature in culture for 3-4 weeks with half-media changes every alternate day. DOX and Ara-C were withdrawn after D3. All experiments were performed between D21-D30.

### CRISPR-editing to generate SNCA-KO iPSC line

The online tool CRISPOR (https://crispor.gi.ucsc.edu/) was used to generate guides targeting Exon 2 of human SNCA^72^. Sense and antisense oligonucleotides encoding SNCA_G4_FOR (GTGCTGCTGAGAAAACCAAAC) and SNCA_G4_REV (GTTTGGTTTTCTCAGCAGCAC) were designed with 5’ CACC- and 5’ AAAC-overhangs respectively, from Sigma. Oligos were annealed using a protocol with a slow gradient drop in temperature from 95°C to 25°C. The annealed oligos were ligated into pSpCas9(BB)-2A-GFP (PX458) vector (Addgene #48138) digested with BbsI restriction enzyme, followed by transformation into chemically competent NEB Stable E.coli (NEB, C3040H). pSpCas9(BB)- 2A-GFP (PX458) was a gift from Feng Zhang (Addgene plasmid # 48138; http://n2t.net/addgene:48138; RRID:Addgene_48138)^73^. Colonies with the correct incorporation of the annealed guides into the vector (PX458-SNCA-G4-GFP) were identified through Sanger sequencing.

iPSCs were plated on 6 well plates at a density of 1 million cells per well in Stemflex media (Gibco, A3349401), supplemented with a CEPT cocktail composed of: Chroman 1 50 nM (Tocris, 7163), Emricasan 5 μM (Tocris, 7310/5), Polyamine supplement 1X (Sigma, P8483-5ML) and Trans-ISRIB 0.7 μM (Tocris, 5284/10)^74^. Four hours following plating, the cells were transfected with 2 μg of PX458-SNCA-G4-GFP plasmid using the Lipofectamine STEM transfection reagent (Invitrogen, STEM00001). Three days following transfection, the GFP +ve cells were sorted as single cells into 96 well plates through FACS using a BD Influx Cell Sorter (BD Biosciences, USA). The plates were incubated for ten days to allow for clonal expansion from single cells, with media changes every three days. The iPSC colonies were then collected for sequence analysis and subsequent expansion. PCR reactions were setup to amplify a 700 kb fragment surrounding the expected edit site at Exon 2 of the SNCA gene and submitted for Sanger sequencing. iPSC clones with a homozygous deletion were selected for downstream analysis (SNCA-KO). WT and SNCA-KO iPSCs were characterized by immunocytochemistry for pluripotency markers (Supplementary Fig. 1a). Successful knockout of SNCA in the selected clone was validated by western blotting and immunocytochemistry of neurons 21 days after differentiation (Supplementary Fig. 1 b/c.).

### Preparation of lentiviral constructs

A third generation lentiviral FUGW vector (Addgene #197034) was used as a vector backbone to generate all lentiviral constructs used in this study. aSYN variants tagged to mScarlet-I and Synaptophysin-mEmerald were cloned downstream of the human Synapsin promoter using Gibson assembly (NEB, E2621S). Synaptophsyin fragment was amplified from cDNA derived from SH-SY5Y cells. mScarlet-I was derived from pmScarlet_Giantin_C1 (Addgene # 85048), mEmerald was derived from mEmerald-Sec61b-C1 (Addgene # 90992). The assembled vectors were verified through sequencing and transfected into HEK293FT cells along with helper plasmids pMDLg/pRRE (Addgene #12251), pRSV-Rev (Addgene #12253), and pMD2.G (Addgene #12259) for viral packaging. Lipofectamine 3000 (Invitrogen, L3000008) was used for transfection according to the manufacturer’s protocol. The following day, the Lipofectamine-containing media was replaced with fresh media supplemented with ViralBoost Reagent (Alstem, VB100) at 2 μL/mL. Supernatants containing viral particles were collected 72 hours after transfection and concentrated using Lenti-X™ Concentrator (Takara Bio, 631231). Viral titers were estimated with a Lentivirus qPCR Quantification Kit (Abcam, ab289841). iPSC-derived neurons were transduced with lentivirus at MOI 3 for aSYN-mScarlet variants and MOI 6 for Synaptophysin-mEmerald between D20-D24 after starting neuronal differentiation and incubated for five days before proceeding to confocal microscopy. For western blotting experiments, aSYN-mScarlet variants were transduced at MOI 1 and incubated for five days before lysis. FSW_jGCaMP8f was a gift from Ronald Hart (Addgene plasmid # 197034; http://n2t.net/addgene:197034; RRID:Addgene_197034). pmScarlet_Giantin_C1 was a gift from Dorus Gadella (Addgene plasmid # 85048; http://n2t.net/addgene:85048; RRID:Addgene_85048)^65^. mEmerald-Sec61b-C1 was a gift from Jennifer Lippincott-Schwartz (Addgene plasmid # 90992; http://n2t.net/addgene:90992; RRID:Addgene_90992)^75^. Plasmids pMDLg/pRRE (Addgene #12251; http://n2t.net/addgene:12251; RRID:Addgene_12251)^76^, pRSV-Rev (Addgene #12253; http://n2t.net/addgene:12253; RRID:Addgene_12253)^76^, and pMD2.G (Addgene #12259 encoding VSV-G; http://n2t.net/addgene:12259; RRID:Addgene_12259) were gifts from Didier Trono.

### Confocal microscopy iPSC-derived neurons

iPSC-derived neurons transduced with aSYN-mScarlet and Synaptophysin-mEmerald were imaged live on an LSM 980 confocal microscope (Carl Zeiss Ltd, UK) using a 40X oil objective. mScarlet and mEmerald were excited with the 561 nm and 488 nm lasers, respectively. Images were obtained at 2x Nyquist sampling, 2x zoom and processed post-acquisition using the LSM Plus (deconvolution) processing feature to generate 16 bit images. For experiments using 1,6 hexanediol, a stock solution of 6% 1,6 hexanediol was prepared in complete media and added to the neurons to reach a final concentration of 3%. Images were obtained at 2 and 4 minutes post treatment, after which hexanediol containing media was replaced with fresh media to monitor recovery of synaptic enrichment at 2 and 4 minutes. FRAP assays on aSYN-mScarlet was performed over a time course of 250 cycles using a 40X objective at 6x zoom, 128 x 128 pixel resolution, with a frame time of 117.9 ms. The 561 nm laser was used for bleaching at 100% laser intensity over 25 iterations. Quantification of fluorescence recovery kinetics was performed using the FRAP profiler plugin on FIJI^66^.

### Quantification and statistical analysis

Data analysis and statistical analysis was performed using Excel 2016 and GraphPad Prism 10.4.1. All data are represented as mean ± standard error (s.d.) if not stated otherwise. Statistical analysis was carried out using unpaired two-tailed t-test, one-way ANOVA with Dunnett’s multiple comparison test, or two-way ANOVA with Tukey’s multiple comparisons tests, respectively as indicated. For 1,6-hexanediol experiments RM one-way ANOVA with Geisser-Greenhouse correction and Dunnett’s multiple comparisons test were performed. Statistical parameters are reported in the Figures and the corresponding Figure Legends. Exact p-values are shown throughout the manuscript. Data distribution was assumed to be normal but this was not formally tested. No statistical methods were used to pre-determine sample sizes but our sample sizes are similar to those reported in previous publications^40,46,77^. Samples were randomly allocated into experimental groups. Data collection and analysis have been performed blinded when indicated. Data were included if the control (wild-type) showed appropriate condensate formation.

## Data Availability

Source data are provided with this study. Plasmids generated in this study are available from the corresponding author with a completed material transfer agreement. All other data supporting the findings of this study are available from the corresponding author on reasonable request.

## Acknowledgements

J.L. acknowledges funding from the Royal Society (Royal Society Dorothy Hodgkin Research Fellowship, DHF/R1/201228), the Addenbrooke’s Charitable Trust (grant award 900325, grant award 900427), the Leverhulme Trust (research project grant, RPG-2022-257) and a Career Support Fund from the University of Cambridge. Furthermore, we acknowledge microscope equipment funding (LSM980 Alpha/Airyscan2, MRC funded, MR/Y002172/1; LSM980 Omega, Clinical School and CIMR Core Facility funded). T.F.O. is funded by DFG (MBExC and SFB1286, project B8). L.A. was funded by Fundação para a Ciência e Tecnologia (2020.05944.BD). Additionally, J.L. and A.C. thank M. Gammons for her advice on CRISPR/Cas9 editing.

## Author contributions

Conceptualization was the responsibility of J.L. Methodology was the responsibility of A.C., A.A., L.A., T.F.O. and J.L. Investigation was the responsibility of A.C., A.A., T.W., L.A, T.F.O. and J.L. Writing of the original draft was conducted by J.L. Review and editing was carried out by A.C., A.A., T.F.O and J.L. Funding acquisition was the responsibility of T.F.O. and J.L. Supervision was carried out by S.R.C., T.F.O. and J.L.

---

**Extended Data Figure 1.**
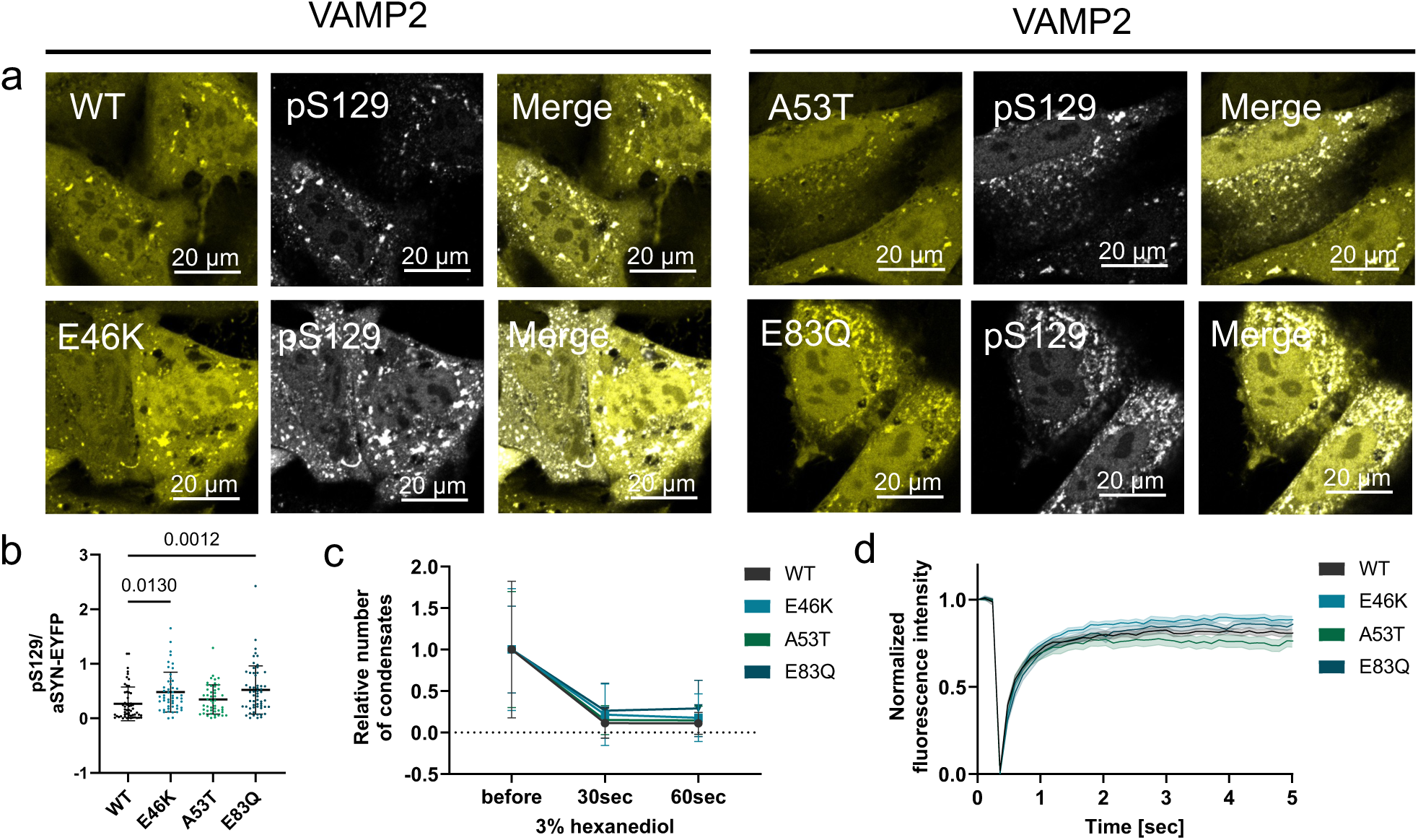
a, Immunocytochemistry for aSYN Ser129 phosphorylation (pS129) in HeLa cells co-expressing VAMP2 with WT aSYN-YFP and other condensate forming aSYN variants. pS129 immunoreactivity is seen in condensates, with increased pS129 intensity for E46K and E83Q. b, Quantification of pS129 staining in a. Data are represented as ratio of pS129 intensity normalized to aSYN-YFP intensity within condensates. n indicates number of cells from 4 biological repeats, n= 55, 50, 48, 58 cells for WT, E46K, A53T and E83Q respectively. Data are shown as mean ± s.d. One-way ANOVA with Dunnett’s multiple comparison test. c, Quantification of aSYN condensate dispersal in HeLa cells upon incubation with 3% 1,6-hexanediol showing a decrease in the relative number of condensates per cell. Data are shown as mean ± s.d. n indicates number of cells from a minimum of 3 biological repeats with n=12, 19, 26, 7 cells for WT, E46K, A53T and E83Q respectively. RM one-way ANOVA with Geisser-Greenhouse correction and Dunnett’s multiple comparisons test. d, Fluorescence recovery after photobleaching (FRAP) experiments demonstrating aSYN recovery for condensates from all condensate forming aSYN-YFP variants. Data are shown as mean ± s.e.m. from a minimum of 4 biological repeats with n=61 for WT, n=36 for E46K, n=41 for A53T, and n=31 for E83Q, with n representing number of condensates measured.

**Extended Data Figure 2.**
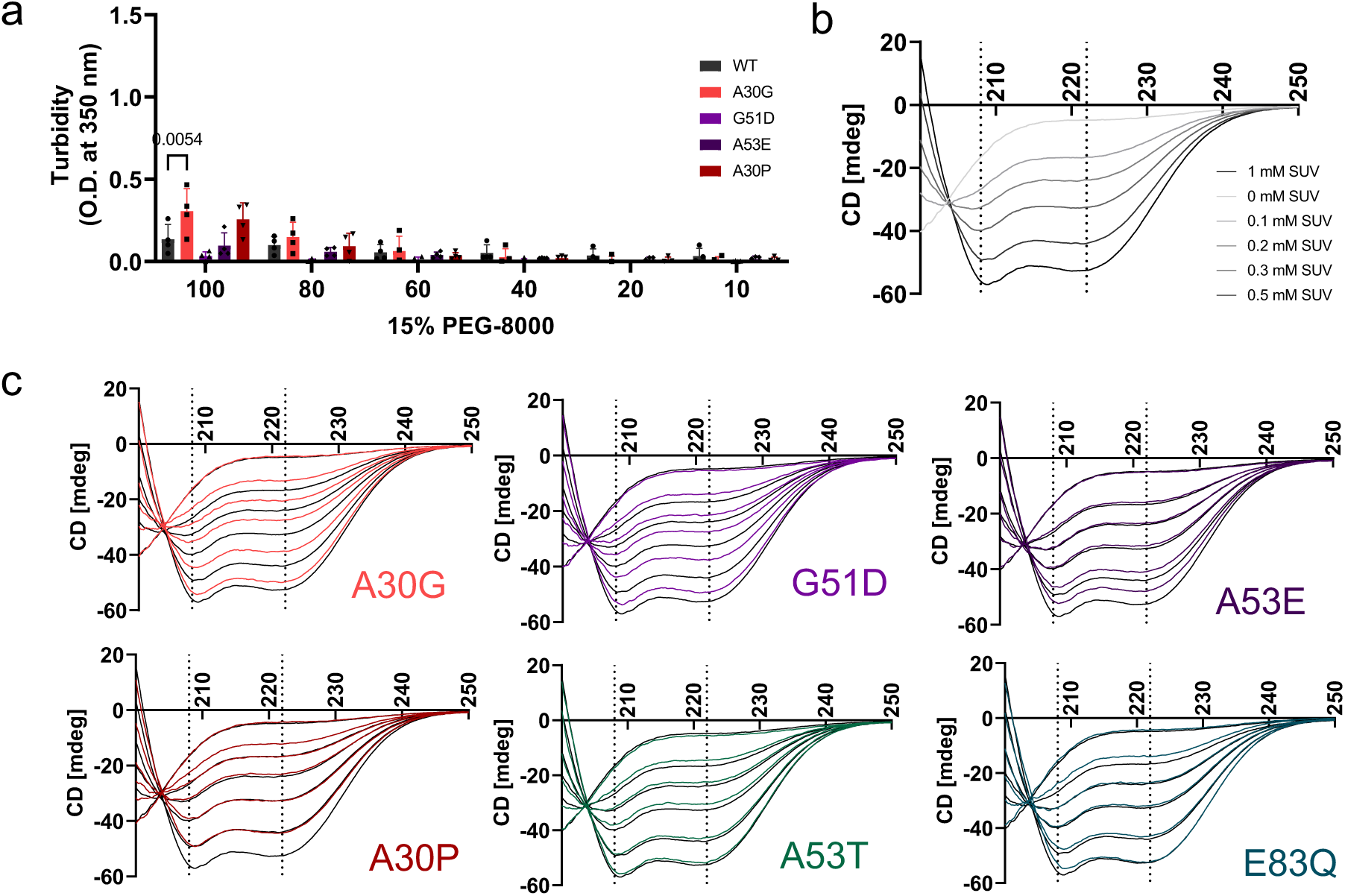
a, Quantification of turbidity data in Fig. 2d. n represent 4 independent repeats from two aSYN batches. Data are shown as mean ± s.d. Two-way ANOVA with Tukey’s multiple comparison test. b, Circular dichroism spectroscopy for WT aSYN without and in the presence of increasing concentrations of small unilamellar vesicles (SUVs), showing transition from random coil conformation to increasing α-helical content. c, Direct comparison of different aSYN variants with WT aSYN showing decreased aSYN α-helical content for A30G, G51D, A53E and A30P. A30P shows the highest reduction in alpha-helical content compared to the other aSYN variants. A53T and E83Q show no difference to WT. All SUVs containing 100% POPS. Data represent at least 4 independent repeats from two aSYN batches.

**Extended Data Figure 3.**
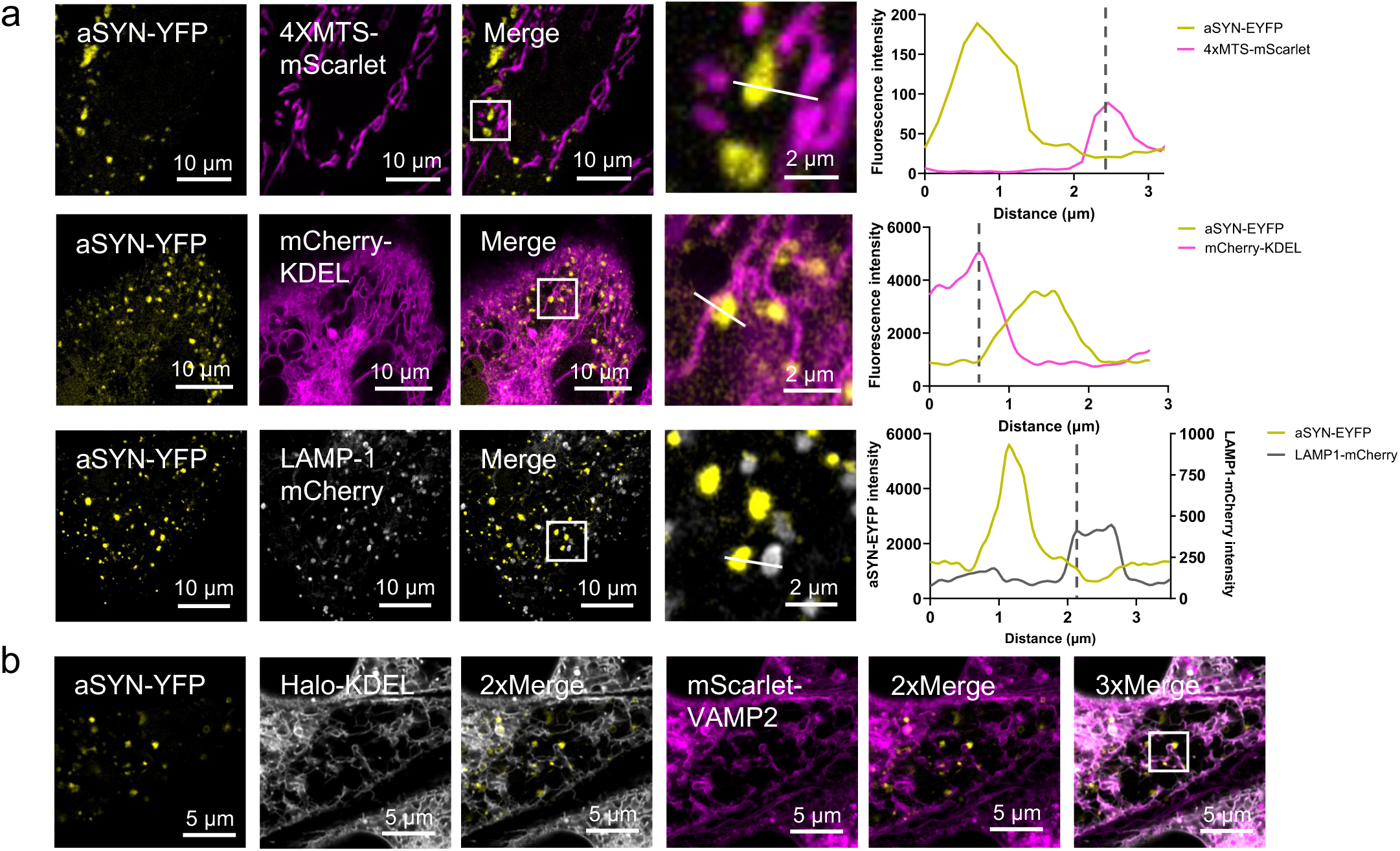
a, HeLa cells co-expressing aSYN-YFP, VAMP2 FL, and organelle markers 4X-MTS-mScarlet, mCherry-KDEL or LAMP1-mCherry for visualization of aSYN condensates on mitochondria, ER or lysosomes, respectively. Representative line diagrams of aSYN condensates and the corresponding organelles, indicating close association of aSYN condensates with the ER membrane b, Three-colour experiment in HeLa cells co-expressing aSYN-YFP, Halo-KDEL and mScarlet-VAMP2 FL showing aSYN condensates on VAMP2 positive ER tubules. Also shown are merged images of aSYN-YFP with Halo-KDEL, aSYN-YFP with mScarlet-VAMP2 and a three-colour merge with aSYN-YFP, Halo-KDEL and mScarlet-VAMP2.

**Supplementary Figure 1.**
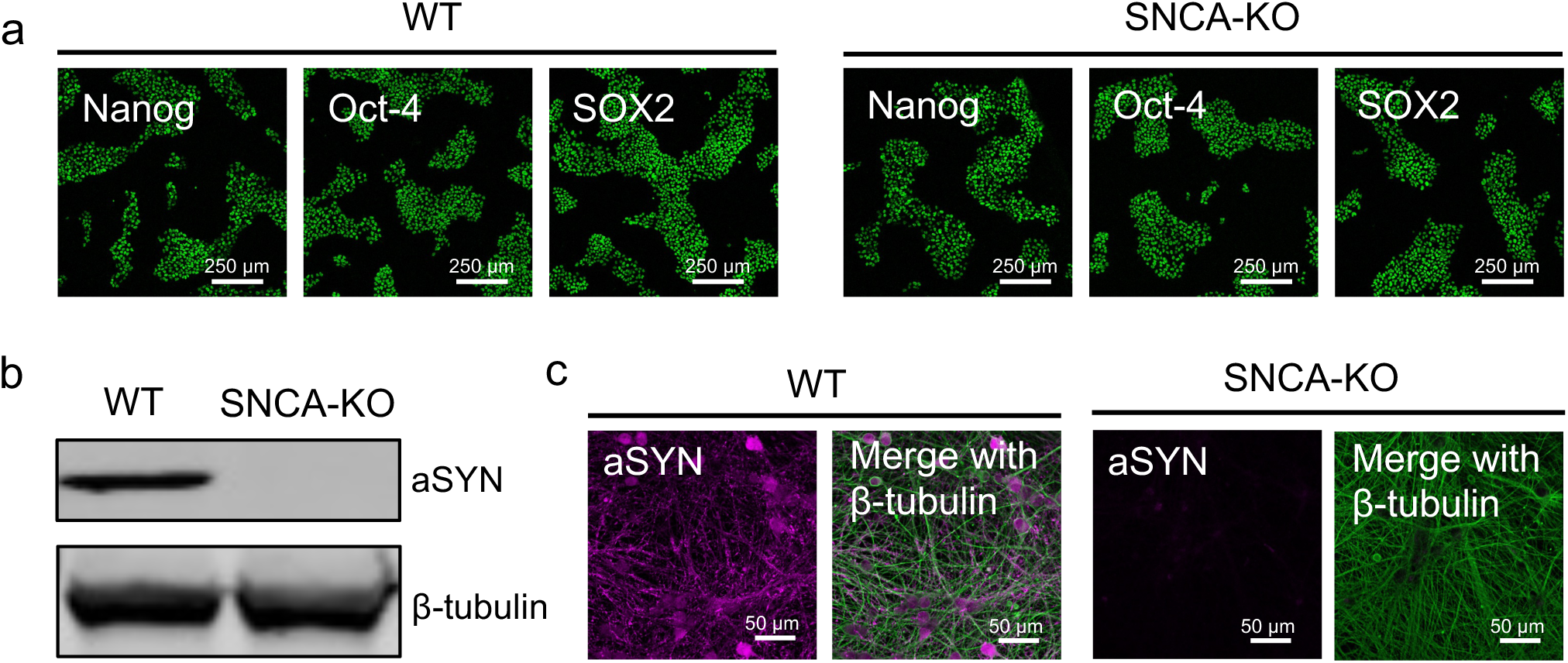
a, Characterisation of WT and SNCA-knock out iPSC lines with stem cell pluripotency markers. b, Western blot showing aSYN expression at D21 for neurons differentiated from WT and SNCA-knock out iPSCs. c, Immunocytochemistry for aSYN and β-tubulin in neurons differentiated from WT and SNCA-knock out iPSCs at D21.

**Extended Data Figure 4.**
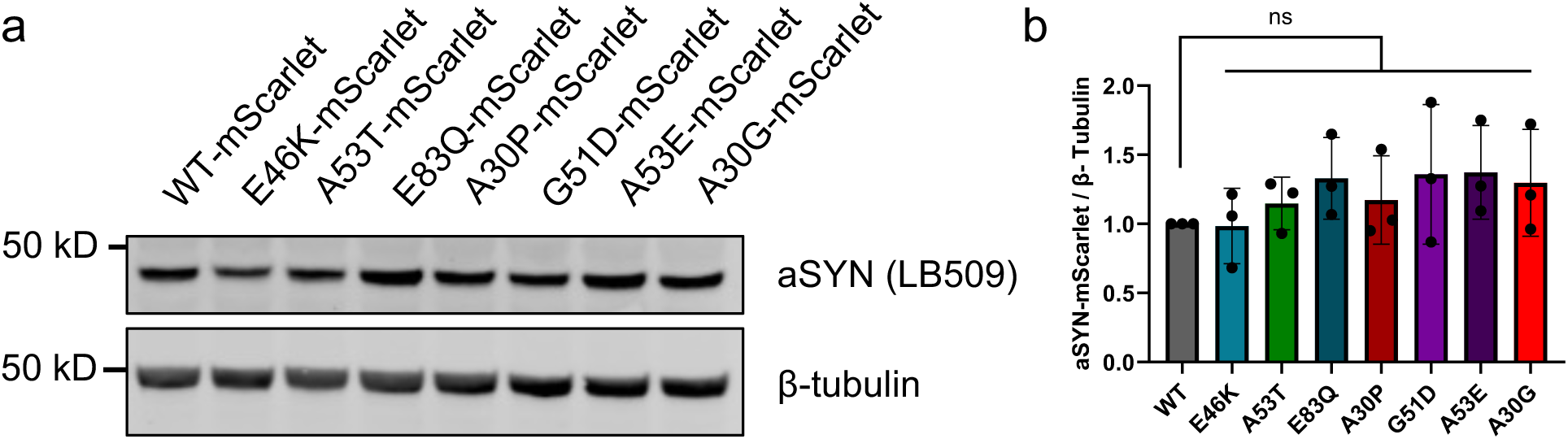
a, aSYN-mScarlet expression levels in iPSC-derived neurons transduced with lentiviruses encoding different aSYN-mScarlet variants between D21-25 and lysed 5 days post transduction. b, Quantification of aSYN levels relative to β-tubulin. n indicates 3 biological repeats. Data are shown as mean ± s.d. One-way ANOVA with Dunnett’s multiple comparison test.

## References

1. Iwai, A. et al. The precursor protein of non-A beta component of Alzheimer’s disease amyloid is a presynaptic protein of the central nervous system. Neuron 14, 467–75 (1995).

2. Maroteaux, L., Campanelli, J. T. & Scheller, R. H. Synuclein: a neuron-specific protein localized to the nucleus and presynaptic nerve terminal. J Neurosci 8, 2804–15 (1988).

3. Spillantini, M. G. et al. α-Synuclein in Lewy bodies. Nature 388, 839–840 (1997).

4. Ray, S. et al. α-Synuclein aggregation nucleates through liquid–liquid phase separation. Nat Chem 12, 705–716 (2020).

5. Hardenberg, M. C. et al. Observation of an α-synuclein liquid droplet state and its maturation into Lewy body-like assemblies. J Mol Cell Biol (2021) doi:10.1093/jmcb/mjaa075.

6. Agarwal, A. et al. VAMP2 regulates phase separation of α-synuclein. Nat Cell Biol 26, 1296–1308 (2024).

7. Wang, C. et al. VAMP2 chaperones α-synuclein in synaptic vesicle co-condensates. Nat Cell Biol 26, 1287–1295 (2024).

8. Mukherjee, S. et al. Liquid-liquid Phase Separation of α-Synuclein: A New Mechanistic Insight for α-Synuclein Aggregation Associated with Parkinson’s Disease Pathogenesis. J Mol Biol 435, 167713 (2023).

9. Zbinden, A., Pérez-Berlanga, M., De Rossi, P. & Polymenidou, M. Phase Separation and Neurodegenerative Diseases: A Disturbance in the Force. Dev Cell 55, 45–68 (2020).

10. Dorsey, E. R. & Bloem, B. R. The Parkinson Pandemic—A Call to Action. JAMA Neurol 75, 9 (2018).

11. Feigin, V. L. et al. Global, regional, and national burden of neurological disorders during 1990–2015: a systematic analysis for the Global Burden of Disease Study 2015. Lancet Neurol 16, 877–897 (2017).

12. Postuma, R. B. et al. Validation of the MDS clinical diagnostic criteria for Parkinson’s disease. Movement Disorders 33, 1601–1608 (2018).

13. Brás, J., Gibbons, E. & Guerreiro, R. Genetics of synucleins in neurodegenerative diseases. Acta Neuropathol 141, 471–490 (2021).

14. Blauwendraat, C., Nalls, M. A. & Singleton, A. B. The genetic architecture of Parkinson’s disease. The Lancet Neurology Preprint at 10.1016/S1474-4422(19)30287-X (2020).

15. Lautenschläger, J., Kaminski, C. F. & Kaminski Schierle, G. S. α-Synuclein – Regulator of Exocytosis, Endocytosis, or Both? Trends Cell Biol 27, 468–479 (2017).

16. Boeynaems, S. et al. Protein Phase Separation: A New Phase in Cell Biology. Trends Cell Biol 28, 420–435 (2018).

17. Kumar, S. T. et al. A NAC domain mutation (E83Q) unlocks the pathogenicity of human alpha-synuclein and recapitulates its pathological diversity. Sci Adv 8, (2022).

18. Anderson, J. P. et al. Phosphorylation of Ser-129 is the dominant pathological modification of α-synuclein in familial and sporadic lewy body disease. Journal of Biological Chemistry (2006) doi:10.1074/jbc.M600933200.

19. Fujiwara, H. et al. α-synuclein is phosphorylated in synucleinopathy lesions. Nat Cell Biol (2002) doi:10.1038/ncb748.

20. Volpicelli-Daley, L. A. et al. Exogenous α-Synuclein Fibrils Induce Lewy Body Pathology Leading to Synaptic Dysfunction and Neuron Death. Neuron (2011) doi:10.1016/j.neuron.2011.08.033.

21. Li, J., Uversky, V. N. & Fink, A. L. Effect of Familial Parkinson’s Disease Point Mutations A30P and A53T on the Structural Properties, Aggregation, and Fibrillation of Human α-Synuclein. Biochemistry 40, 11604–11613 (2001).

22. Mohite, G. M. et al. Comparison of Kinetics, Toxicity, Oligomer Formation, and Membrane Binding Capacity of α-Synuclein Familial Mutations at the A53 Site, Including the Newly Discovered A53V Mutation. Biochemistry 57, 5183–5187 (2018).

23. Stephens, A. D. et al. Extent of N-terminus exposure of monomeric alpha-synuclein determines its aggregation propensity. Nat Commun 11, 2820 (2020).

24. Alberti, S., Gladfelter, A. & Mittag, T. Considerations and Challenges in Studying Liquid-Liquid Phase Separation and Biomolecular Condensates. Cell 176, 419–434 (2019).

25. Jo, E., Fuller, N., Rand, R. P., St George-Hyslop, P. & Fraser, P. E. Defective membrane interactions of familial Parkinson’s disease mutant A30P α-synuclein 1 1Edited by I. B. Holland. J Mol Biol 315, 799–807 (2002).

26. Outeiro, T. F. & Lindquist, S. Yeast cells provide insight into alpha-synuclein biology and pathobiology. Science 302, 1772–5 (2003).

27. Wang, C. et al. Scalable Production of iPSC-Derived Human Neurons to Identify Tau-Lowering Compounds by High-Content Screening. Stem Cell Reports 9, 1221–1233 (2017).

28. Fernandopulle, M. S. et al. Transcription Factor-Mediated Differentiation of Human iPSCs into Neurons. Curr Protoc Cell Biol 79, e51 (2018).

29. Imoto, Y. et al. Dynamin is primed at endocytic sites for ultrafast endocytosis. Neuron 110, 2815–2835.e13 (2022).

30. Yoshida, T. et al. Compartmentalization of soluble endocytic proteins in synaptic vesicle clusters by phase separation. iScience 26, 106826 (2023).

31. Xu, B. et al. Distinct Effects of Familial Parkinson’s Disease-Associated Mutations on α-Synuclein Phase Separation and Amyloid Aggregation. Biomolecules 13, (2023).

32. Kapasi, A. et al. A novel SNCA E83Q mutation in a case of dementia with Lewy bodies and atypical frontotemporal lobar degeneration. Neuropathology 40, 620–626 (2020).

33. Conway, K. A., Harper, J. D. & Lansbury, P. T. Accelerated in vitro fibril formation by a mutant alpha-synuclein linked to early-onset Parkinson disease. Nat Med 4, 1318–20 (1998).

34. Parra-Rivas, L. A. et al. Serine-129 phosphorylation of α-synuclein is an activity-dependent trigger for physiologic protein-protein interactions and synaptic function. Neuron 111, 4006–4023.e10 (2023).

35. Ramalingam, N. et al. Dynamic physiological α-synuclein S129 phosphorylation is driven by neuronal activity. NPJ Parkinsons Dis 9, 4 (2023).

36. Ramalingam, N., Brontesi, L., Jin, S., Selkoe, D. J. & Dettmer, U. Dynamic reversibility of α-synuclein serine-129 phosphorylation is impaired in synucleinopathy models. EMBO Rep 24, (2023).

37. Ray, S. et al. Divergent effects of pathological α-synuclein truncations and mutations on phase separation. bioRXIV Preprint at 10.1101/2024.11.18.624073 (2024).

38. Middleton, E. R. & Rhoades, E. Effects of Curvature and Composition on α-Synuclein Binding to Lipid Vesicles. Biophys J 99, 2279–2288 (2010).

39. Bodner, C. R., Maltsev, A. S., Dobson, C. M. & Bax, A. Differential Phospholipid Binding of α-Synuclein Variants Implicated in Parkinson’s Disease Revealed by Solution NMR Spectroscopy. Biochemistry 49, 862–871 (2010).

40. Wu, X. et al. RIM and RIM-BP Form Presynaptic Active-Zone-like Condensates via Phase Separation. Mol Cell 73, 971–984.e5 (2019).

41. Snead, W. T. et al. Membrane surfaces regulate assembly of ribonucleoprotein condensates. Nat Cell Biol 24, 461–470 (2022).

42. Gao, Y. et al. Lipid-mediated phase separation of AGO proteins on the ER controls nascent-peptide ubiquitination. Mol Cell 82, 1313–1328.e8 (2022).

43. Fares, M.-B. et al. The novel Parkinson’s disease linked mutation G51D attenuates in vitro aggregation and membrane binding of -synuclein, and enhances its secretion and nuclear localization in cells. Hum Mol Genet 23, 4491–4509 (2014).

44. Ysselstein, D. et al. Effects of impaired membrane interactions on α-synuclein aggregation and neurotoxicity. Neurobiol Dis (2015) doi:10.1016/j.nbd.2015.04.007.

45. Ghosh, D. et al. The Newly Discovered Parkinson’s Disease Associated Finnish Mutation (A53E) Attenuates α-Synuclein Aggregation and Membrane Binding. Biochemistry 53, 6419–6421 (2014).

46. Park, D. et al. Cooperative function of synaptophysin and synapsin in the generation of synaptic vesicle-like clusters in non-neuronal cells. Nat Commun 12, 263 (2021).

47. Fortin, D. L. et al. Lipid Rafts Mediate the Synaptic Localization of α-Synuclein. The Journal of Neuroscience 24, 6715–6723 (2004).

48. West, S. et al. G51D mutation of the endogenous rat *Snca* gene disrupts synaptic localisation of α-synuclein priming for Lewy-like pathology. BioRXIV Preprint at 10.1101/2023.10.27.564027 (2023).

49. Piroska, L. et al. α-Synuclein liquid condensates fuel fibrillar α-synuclein growth. Sci Adv 9, eadg5663 (2023).

50. Fusco, G. et al. Structural basis of membrane disruption and cellular toxicity by α-synuclein oligomers. Science (1979) 358, 1440–1443 (2017).

51. Mohite, G. M. et al. The Familial α-Synuclein A53E Mutation Enhances Cell Death in Response to Environmental Toxins Due to a Larger Population of Oligomers. Biochemistry 57, 5014–5028 (2018).

52. Stefanovic, A. N. D., Lindhoud, S., Semerdzhiev, S. A., Claessens, M. M. A. E. & Subramaniam, V. Oligomers of Parkinson’s Disease-Related α-Synuclein Mutants Have Similar Structures but Distinctive Membrane Permeabilization Properties. Biochemistry 54, 3142–50 (2015).

53. Conway, K. A. et al. Acceleration of oligomerization, not fibrillization, is a shared property of both α-synuclein mutations linked to early-onset Parkinson’s disease: Implications for pathogenesis and therapy. Proc Natl Acad Sci U S A (2000) doi:10.1073/pnas.97.2.571.

54. Xu, C. et al. The Pathological G51D Mutation in Alpha-Synuclein Oligomers Confers Distinct Structural Attributes and Cellular Toxicity. Molecules 27, 1293 (2022).

55. Martikainen, M. H., Päivärinta, M., Hietala, M. & Kaasinen, V. Clinical and imaging findings in Parkinson disease associated with the A53E SNCA mutation. Neurol Genet 1, e27 (2015).

56. Pasanen, P. et al. SNCA mutation p.Ala53Glu is derived from a common founder in the Finnish population. Neurobiol Aging 50, 168.e5–168.e8 (2017).

57. Pasanen, P. et al. Novel α-synuclein mutation A53E associated with atypical multiple system atrophy and Parkinson’s disease-type pathology. Neurobiol Aging 35, 2180.e1–5 (2014).

58. Kiely, A. P. et al. α-Synucleinopathy associated with G51D SNCA mutation: a link between Parkinson’s disease and multiple system atrophy? Acta Neuropathol 125, 753–69 (2013).

59. Lesage, S. et al. G51D α-synuclein mutation causes a novel parkinsonian-pyramidal syndrome. Ann Neurol 73, 459–71 (2013).

60. Tokutake, T. et al. Clinical and neuroimaging features of patient with early-onset Parkinson’s disease with dementia carrying SNCA p.G51D mutation. Parkinsonism Relat Disord 20, 262–4 (2014).

## References

61. Phua, S. C. et al. Dynamic Remodeling of Membrane Composition Drives Cell Cycle through Primary Cilia Excision. Cell 168, 264–279.e15 (2017).

62. Chertkova, A. O. et al. Robust and Bright Genetically Encoded Fluorescent Markers for Highlighting Structures and Compartments in Mammalian Cells. bioRXIV Preprint at 10.1101/160374 (2017).

63. Van Engelenburg, S. B. & Palmer, A. E. Imaging type-III secretion reveals dynamics and spatial segregation of Salmonella effectors. Nat Methods 7, 325–30 (2010).

64. Campbell, R. E. et al. A monomeric red fluorescent protein. Proc Natl Acad Sci U S A 99, 7877–82 (2002).

65. Bindels, D. S. et al. mScarlet: a bright monomeric red fluorescent protein for cellular imaging. Nat Methods 14, 53–56 (2017).

66. Schindelin, J., et al. Fiji: an open-source platform for biological-image analysis. Nat Methods 9, 676–82 (2012).

67. Man, W. K. et al. The docking of synaptic vesicles on the presynaptic membrane induced by α-synuclein is modulated by lipid composition. Nat Commun 12, 927 (2021).

68. Modzelewski, A. J. et al. A mouse-specific retrotransposon drives a conserved Cdk2ap1 isoform essential for development. Cell 184, 5541–5558.e22 (2021).

69. Chen, Y. H., Yang, J. T. & Martinez, H. M. Determination of the secondary structures of proteins by circular dichroism and optical rotatory dispersion. Biochemistry 11, 4120– 31 (1972).

70. Weinberger, A. et al. Gel-assisted formation of giant unilamellar vesicles. Biophys J 105, 154–64 (2013).

71. Gietz, R. D. & Schiestl, R. H. High-efficiency yeast transformation using the LiAc/SS carrier DNA/PEG method. Nat Protoc 2, 31–34 (2007).

72. Concordet, J.-P. & Haeussler, M. CRISPOR: intuitive guide selection for CRISPR/Cas9 genome editing experiments and screens. Nucleic Acids Res 46, W242–W245 (2018).

73. Ran, F. A. et al. Genome engineering using the CRISPR-Cas9 system. Nat Protoc 8, 2281–2308 (2013).

74. Chen, Y. et al. A versatile polypharmacology platform promotes cytoprotection and viability of human pluripotent and differentiated cells. Nat Methods 18, 528–541 (2021).

75. Nixon-Abell, J. et al. Increased spatiotemporal resolution reveals highly dynamic dense tubular matrices in the peripheral ER. Science 354, (2016).

76. Dull, T. et al. A third-generation lentivirus vector with a conditional packaging system. J Virol 72, 8463–71 (1998).

77. Park, D. et al. Synaptic vesicle proteins and ATG9A self-organize in distinct vesicle phases within synapsin condensates. Nat Commun 14, 455 (2023).

