## Supplemental Table 1 for "Dual effect of α-synuclein disease variants on condensate formation"

Supplementary Table 1. List of antibodies used in this study.

| **Antibody** | **Catalog No.** | **Application** | **Dilution** |
| --- | --- | --- | --- |
| **Primary** | | | |
| Mouse Purified anti-α-Synuclein Phospho (Ser129) Antibody, clone 81A | BioLegend, 825701 | ICC | 1:1000 |
| StemLight Pluripotency Transcription Factor Antibody Kit (Nanog, Oct4A, SOX2) | CST, 9093 | ICC | 1:1000 |
| Mouse alpha Synuclein antibody | Synaptic Systems, 128 211 | ICC | 1:250 |
| β3-Tubulin (D65A4) XP® Rabbit mAb | CST, 5666 | ICC | 1:500 |
| Alpha-synuclein Monoclonal Antibody (LB509) | Invitrogen, 180215 | WB | 1:1000 |
| β3-Tubulin (D65A4) XP® Rabbit mAb | CST, 5666 | WB | 1:1000 |
| **Secondary** | | | |
| F(ab')2-Goat anti-Mouse IgG (H+L) Cross-Adsorbed Secondary Antibody, Alexa Fluor™ 594 | Invitrogen, A11020 | ICC | 1:1000 |
| F(ab')2-Goat anti-Rabbit IgG (H+L) Cross-Adsorbed Secondary Antibody, Alexa Fluor™ 488 | Invitrogen, A11070 | ICC | 1:1000 |
| F(ab')2-Goat anti-Mouse IgG (H+L) Cross-Adsorbed Secondary Antibody, Alexa Fluor™ 488 | Invitrogen, A11017 | ICC | 1:1000 |
| F(ab')2-Goat anti-Rabbit IgG (H+L) Cross-Adsorbed Secondary Antibody, Alexa Fluor™ 594 | Invitrogen, A11072 | ICC | 1:1000 |
| Goat anti-Mouse IgG (H+L) Secondary Antibody, DyLight™ 800 | Invitrogen, SA535521 | WB | 1:10000 |
| Goat anti-Rabbit IgG (H+L) Secondary Antibody, DyLight™ 680 | Invitrogen, 35568 | WB | 1:10000 |
