## Supplemental Table 2 for "Dual effect of α-synuclein disease variants on condensate formation"

Supplementary Table 2. List of yeast strains and plasmids used in this study.

| Strain | Genotype | Source |
| --- | --- | --- |
| BY4741 | *MATa his3Δ1 leu2Δ0 met15Δ0 ura3Δ0* | Euroscarf |
| BY4741 pGPD-GFP | BY4741 harboring *p426GPDpr-GFP* | This Study |
| BY4741 pGPD-SNCA (WT) - GFP | BY4741 harboring  *p426GPDpr*-SNCA (WT)-GFP | 10.1126/science.1090439. |
| BY4741 pGPD-SNCA (A53T)-GFP | BY4741 harboring  *p426GPDpr*-SNCA (A53T)-GFP | 10.1126/science.1090439. |
| BY4741 pGPD-SNCA (A53E)-GFP | BY4741 harboring  *p426GPDpr*-SNCA (A53E)-GFP | This Study |
| BY4741 pGPD-SNCA (E46K)-GFP | BY4741 harboring  *p426GPDpr*-SNCA (E46K)-GFP | This Study |
| BY4741 pGPD-SNCA (A30G)-GFP | BY4741 harboring  *p426GPDpr*-SNCA (A30G)-GFP | This Study |
| BY4741 pGPD-SNCA (A30P)-GFP | BY4741 harboring  *p426GPDpr*-SNCA (A30P)-GFP | This Study |
| BY4741 pGPD-SNCA (G51D)-GFP | BY4741 harboring  *p426GPDpr*-SNCA G51D)-GFP | This Study |
